# Transcriptome Assembly at Single-Cell Resolution with Beaver

**DOI:** 10.1101/2024.11.04.621958

**Authors:** Qian Shi, Qimin Zhang, Mingfu Shao

## Abstract

Emerging single-cell RNA sequencing techniques (scRNA-seq) has enabled the study of cellular transcriptome heterogeneity, yet accurate reconstruction of full-length transcripts at single-cell resolution remains challenging due to high dropout rates and sparse coverage. While meta-assembly approaches offer promising solutions by integrating information across multiple cells, current methods struggle to balance consensus assembly with cell-specific transcriptional signatures. Here, we present Beaver, a cell-specific transcript assembler designed for short-read scRNA-seq data. Beaver implements a transcript fragment graph to organize individual assemblies and designs an efficient dynamic programming algorithm that searches for candidate full-length transcripts from the graph. Beaver incorporates two random forest models trained on 51 meticulously engineered features that accurately estimate the likelihood of each candidate transcript being expressed in individual cells. Our experiments, performed using both real and simulated Smart-seq3 scRNA-seq data, firmly show that Beaver substantially outperforms existing meta-assemblers and single-sample assemblers. At the same level of sensitivity, Beaver achieved 32.0%-64.6%, 13.5%-36.6%, and 9.8%-36.3% higher precision in average compared to meta-assemblers Aletsch, TransMeta, and PsiCLASS, respectively, with similar improvements over single-sample assemblers Scallop2 (10.1%-43.6%) and StringTie2 (24.3%-67.0%). Beaver is freely available at https://github.com/Shao-Group/beaver. Scripts that reproduce the experimental results of this manuscript are available at https://github.com/Shao-Group/beaver-test.

## 1 Introduction

The landscape of cellular transcriptomes remained largely unexplored until the emergence of single-cell sequencing technologies, which expanded our understanding of cellular heterogeneity and transcriptional dynamics. This technological breakthrough has evolved from basic gene expression profiling to analyses capable of detecting transcript isoforms, allele-specific expression, and complex regulatory patterns at the individual cell level. Unlike traditional bulk RNA sequencing methods which provided averaged expression profiles across cell populations, scRNA-seq has unveiled cell-to-cell variations, enabling identification of rare cell types and transitional states. Among the various scRNA-seq protocols, droplet-based platforms such as 10X Genomics Chromium [34,12] excel in throughput, and full-length transcript sequencing methods like Smart-seq series [18,4,3] offer deeper insights into transcript architecture and splice variants [28].

The majority of scRNA-seq analyses remain focusing on gene-level expression [5,24], rather than the rich diversity of transcript isoforms. High dropout rates, sparse coverage, and PCR amplification bias collectively complicate accurate isoform detection and quantification at the cellular level [30]. On the other hand, transcript assembly, the computational reconstruction of full-length transcripts from sequencing reads, has been extensively developed for bulk RNA-seq data, including Cufflinks [27], CLASS2 [22], the StringTie series [17,7], and the Scallop series [19,33], to name just a few. However, the direct application of these bulk RNA-seq assemblers to single-cell data has proven challenging due to data sparsity, where dropout events and coverage gaps can result in fragmented assemblies. These single-sample assemblers, while naturally maintaining cell specificity, often yield limited full-length transcripts by overlooking shared information across cells. This has motivated the development of specialized single-cell assemblers, such as scRNAss [11] and RNA-Bloom [14], but their performances are limited by the guidance of known transcriptome references.

It is much desirable for an assembly method to utilize the share information from multiple cells to recover full-length transcripts while preserving cell-specific expression landscapes. Meta-assembly, which reconstructs expressed transcripts from multiple samples, offers a promising direction. Several dedicated algorithms have been developed in this field [1,10,25,13], including PsiCLASS [23], TransMeta [31], and Aletsch [20]. However, limitations persist in applying existing meta-assemblers to single-cell data. The primary objective of meta-assembly is seeking consensus across samples. Single-cell analysis, on the contrary, demands cell-specific assemblies that retain individual transcriptional signatures. Current meta-assemblers developed distinct strategies to balance global consensus with sample-specific accuracy. PsiCLASS [23] achieves this balance through a voting mechanism, but in practice, its performance deteriorates with low-coverage samples—a common scenario of single-cell data. TransMeta [31] prioritizes meta-assembly accuracy by constructing a combined graph from all input alignments, then distributing transcripts to individual samples based on junction coverage thresholds. This strategy often fails to preserve cell-specific characteristics due to indiscriminate junction sharing. The most recent tool, Aletsch [20], introduces a hybrid approach by constructing both combined and individual cell-specific splice graphs. Its conservative strategy prioritizes cell-specific assemblies, but still struggles with transcript fragmentation when multiple exons and splicing junctions are missing.

To address these limitations, we introduce Beaver, a transcript assembler designed to reconstruct accurate cell-specific transcriptomes using short-read scRNA-seq data. Our approach is motivated by the observation that while dropout events create gaps in individual cell assemblies, the missing information often exists in other cells. Beaver follows this biological insight to reconstruct full-length transcripts while carefully preserving cell-specific expression patterns. We introduced a transcript fragment graph that organizes individual assemblies, allowing for reconstructing full-length transcripts from the fragments from different cells. An efficient dynamic programming algorithm selects high-quality candidates in the graph by optimizing a merging score based on junction compatibility and coverage. We engineered 30 features to characterize true isoforms, and 21 cell-specific features to estimate transcript expression likelihood in individual cells. Equipped with these informative features, Beaver trains two random forest models that first conduct coarse-grained filtering and then perform fine-grained cell-specific scoring, achieving accurate assembly at single-cell resolution. Our experimental results show that, on both real Smart-seq3 scRNA-seq data and simulated datasets spanning various cell populations, Beaver drastically outperforms leading meta-assemblers (TransMeta, PsiCLASS, and Aletsch) and single-sample assemblers (StringTie2 and Scallop2).

## 2 Methods

Beaver reconstructs full-length transcripts for individual cells from single-cell RNA sequencing data. It integrates cross-cell information while preserving cell-specific transcriptional characteristics. Beaver’s method consists of four main steps: collection of individual cell assemblies, construction of transcript fragment graphs, full-length transcript generation, and assignment of cell-specific scores.

### 2.1 Collection of Individual Assemblies

Beaver takes as input an assembly (a set of assembled transcripts) for each individual cell. Each transcript *t* is required to be associated with a normalized coverage from 0 to 1, denoted as *score*(*t*), indicating its reliability. These inputs can be generated using any single-cell assembler or meta-assembler that produces individual assemblies. In this study, we select Aletsch [20] as our individual assemblies provider because of its effectiveness in three aspects: generating transcripts with reliable confidence scores, capturing shared information across cells, and maintaining cell-specific characteristics by limiting excessive information integration. Beaver recognizes that many “transcripts” in individual assemblies are actually fragments, which we refer to as “transcript fragments” throughout. Beaver assembles these transcript fragments and applies cell-specific scoring to produce individual assemblies with enhanced accuracy (see Section 3.2).

### 2.2 Transcript Fragment Graph Construction

We construct a directed graph *G* = (*V, E*) to capture connections across transcript fragments from all individual assemblies. Each vertex *v* ∈ *V* represents a transcript fragment, and directed edges *e* = (*u, v*) connect vertices if and only if *u*’s suffix intron-chain overlaps with *v*’s prefix intron-chain (Fig. 1). This edge connection criterion aims for consistency in overlapping regions, allowing conflict-free transcript merging, and also prevents inappropriate junction combination from various transcripts. Single-exon transcripts are excluded from this work, as our primary goal is to extend transcript fragments by identifying missing junctions.

**Fig. 1:**
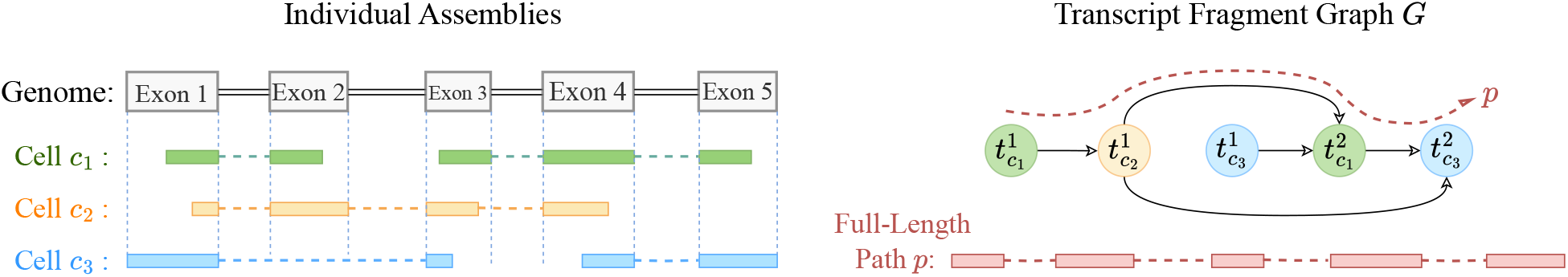
Construction of a transcript fragment graph. Each vertex in *G* represents a transcript fragment *t*_*c*_ from cell *c*, with different colors indicating transcripts from distinct cells. The path *p* ∈ *G* comprising 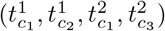 represents a candidate full-length transcript.

The graph *G* consists of multiple connected components, where each component may contain one or more vertices. In practice, many components contains only isolated vertices, indicating transcripts that cannot be merged with others due to unique junction patterns or limited fragmentation in their gene locus. We focus on components with multiple transcript fragments where meaningful merging opportunities exist to extend fragmented chains into full-length transcripts. Within such component, any path through the graph—whether through a single vertex or multiple connected vertices—represents a potential full-length transcript.

### 2.3 Formulation and Algorithm for Path Selection

Let *G*_*c*_ = (*V*_*c*_, *E*_*c*_) be one connected component of *G*. The core challenge of full-length transcript reconstruction lies in identifying the most probable paths through the transcript fragment graph *G* = (*V, E*). We formulate this task as an optimization problem, where the key is designing an objective function that can guide the search for reliable paths. We propose such an objective function, termed *merging-score*, that integrates scores of the given transcript fragments and structural completeness. The intuition behind this merging-score stems from two observations: fragmented transcripts in individual assemblies often require extension to reach the full-length sequences, and true full-length transcripts typically show consistent support across multiple cells.

Let *p* be a path in *G*_*c*_; we also use *p* to represent the corresponding transcript. The merging-score of *p*, denoted as *F* (*p*), is defined as *F* (*p*) := *BJ* (*p*) *· NJ* (*p*), where *BJ* (*p*) is the bottleneck junction-score, defined below, and *NJ* (*p*) is the number of junctions in the corresponding transcript. To define *BJ* (*p*), we first introduce the concept of transcript compatibility. A transcript fragment *t* is compatible with path *p*, denoted as *t* ∼ *p*, if *t* shares at least one junction with *p* and does not contain any conflicting junctions. For a junction *j* ∈ *p*, we define its junction-score, denoted as *J* (*j, p*), as the sum of scores of all transcript fragments that contain junction *j* and are compatible with *p*, i.e., *J* (*j, p*) := ∑_*t*:*j*∈*t* and *t*∼*p*_ *score*(*t*). The bottleneck junction-score of *p* is defined as: *BJ* (*p*) = min_*j*∈*p*_ *J* (*j, p*), i.e., the smallest junction-score among all junctions in path *p*.

We believe this objective function is appropriate for selecting full-length transcripts from the fragment graph. First, by maximizing the bottleneck junction-score, we ensure that selected paths have strong support for all junctions, reducing the likelihood of artificial chimeric transcripts. Second, incorporating the number of junctions into the objective scoring function actively encourages the extension of transcript fragments into full-length transcripts, addressing the fundamental challenge of transcript fragmentation in single-cell RNA-seq data. Finally, our strict compatibility requirement prevents the inappropriate mixing of junctions from arbitrary isoforms, reducing false-positive rates.

#### Dynamic Programming

We design an efficient dynamic programming heuristic to select paths with maximized merging-score. Let 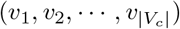 be a topological order of vertices in *G*_*c*_ = (*V*_*c*_, *E*_*c*_). To compute optimal path up to *v*_*j*_, we examine all incoming edges (*v*_*i*_, *v*_*j*_) ∈ *E*_*c*_, and obtain all paths previously computed for each predecessor vertex *v*_*i*_. For computational efficiency, we maintain only the top *p*_*n*_ paths (default: 15) at each vertex using a min-heap structure, where paths are ranked by their merging-scores. Additionally, we limit the total number of paths per connected component to *p*_*c*_ (default: 100), retaining only the highest-scoring candidates. Pseudocode of this heuristic is available at Supplementary Note 1. While these constraints theoretically lead to a suboptimal solution, our empirical testing demonstrates they achieve an effective balance between computational efficiency and transcript reconstruction accuracy. This heuristic successfully identifies promising full-length transcripts, providing a good source of candidates that will undergo more comprehensive evaluation in the subsequent machine learning-based scoring steps.

### 2.4 Scoring Assembled Full-length Transcripts

The above step produces a set of candidate full-length paths/transcripts *P* across all input cells. For each path *p* ∈ *P* and each cell *c*, we now estimate the probability that *p* is expressed in cell *c*, denoted as Pr(*p, c*). We consider only pairs (*p, c*) where at least one junction in *p* gets expressed in cell *c*. To further take into account the computational efficiency, we design an approach that consists of two machine-learning models, namely Beaver-General and Beaver-Specific.

Beaver-General evaluates each candidate full-length transcript *p*, rather than a (path, cell) pair, and produces a confidence score that estimates the likelihood of *p* being correct (regardless of the cells it may be expressed from). Beaver-General serves as a preprocessing step: candidates failing to meet Beaver-General’s score threshold are discarded, yielding a filtered set *P* ^*′*^. Beaver-Specific then estimates Pr(*p, c*) for each *p* ∈ *P* ^*′*^ and every cell, representing the probability that *p* is both correctly assembled and expressed in cell *c*. Both models are implemented as random forests, trained with features detailed below.

#### Feature Engineering

We design two feature sets: 30 “general features” evaluating overall transcript reliability, and 21 “cell-specific” features assessing expression likelihood in specific cells. These features are grouped into 3 categories, with detailed descriptions available in Supplementary Note 2.

1. *Junction Coverage Features* quantify splicing junction support from transcript fragments (in the given individual assemblies) and cells. For each (cell *c*, path *p*, junction *j*) triplet, we calculate junction score similar to the definition in Section 2.3, where support is contributed by transcript fragments *t* that: (1) contain junction *j*, (2) are compatible with path *p*, and (3) belong to cell *c*. To handle varying junction supports across paths, we summarize these coverage values using statistical measures (minimum, median, mean, maximum, and standard deviation).
2. *Cell Support Features* provide global assessment of cellular support for candidate paths. These features include the count of supporting cells (positive factors) and quantitative measures of these factors, such as per-cell coverage levels and supported junction counts.
3. *Fragment Connecting Features* characterize relationships between the input transcript fragments within the merged/candidate full-length transcripts. These features distinguish between transcripts that maintain input intron-chain integrity and those merged from multiple transcript fragments, quantifying fragment contributions and inter-fragment relationships.

#### Training Beaver-General

We implement Beaver-General as a random forest model (n_estimators=100, max_depth=12), using the 30 general features described above. This model evaluates transcripts independently of their cell assignments. Training data comes from chromosomes 1–9, with testing on remaining chromosomes for all datasets (see Section 3.2, 3.3). To label the candidate paths, ground-truth expressed transcripts on chromosomes 1–9 from all cells are unified. A candidate path *p* ∈ *P* is labeled as 1, if the intron-chain of *p* matches one in the unified true expressed transcripts, and 0 otherwise. Beaver-General produces a scores indicating the likelihood of a candidate being correct; candidates scoring below threshold (default: 0.2) are filtered out. This filtering step helps control false positives and ensures balanced samples for the subsequent training of Beaver-Specific.

#### Training Beaver-Specific

Beaver-Specific builds upon the identical random forest configuration. It is trained using all 51 features. This model hence incorporates information from both general transcript characteristics and specific cell-transcript interactions. The training data are also from chromosomes 1–9. Each instance is a (cell, path) pair, labeled 1 only if the path matches an expressed transcript in that cell’s ground truth. The total training samples for Beaver-Specific is much more than Beaver-General’s, as candidate transcripts can appear in multiple cells.

## 3 Results

### 3.1 Experimental Setup

#### Compared Assemblers

We compare Beaver against three leading meta-assemblers TransMeta (v1.0), PsiCLASS (v1.0.3), and Aletsch (v1.1.1), and two popular single-sample assemblers StringTie2 (v2.2.1) and Scallop2 (v1.1.2). All tools were executed with their default parameters. Each tool produce an assembly (a set of assembled transcripts in GTF format) for each individual cell. Beaver takes a prior assembly with transcript coverages as input (rather than reads alignment); in the experiments below, we use the assemblies generated from Aletsch for Beaver.

#### Real Datasets

We conducted experiments on two real Smart-seq3 single-cell RNA-seq datasets (Accession ID E-MTAB-8735): HEK293T, consisting of 192 human kidney epithelial cells, and Mouse-Fibroblast, containing 369 mouse tail fibroblast cells. To ensure robustness across varying cell populations, we analyzed multiple subsets of cells in a wide range of {5, 10, 30, 50, 100, 192} for the HEK293T dataset, and {10, 30, 50, 100, 200, 369} for the Mouse-Fibroblast dataset.

#### Simulated Datasets

Simulated data was generated using the scRNA-seq data simulation pipeline [29] with simulation component of RSEM [9]. We choose RSEM because it learns expression patterns from real RNA-seq data and generates reads based on these learned parameters through its generative model. In this way, the distribution of the simulated reads aligns better with the provided real RNA-seq data (Smart-seq3 scRNA-seq data, in our case). Specifically, for each cell in the above HEK293T and Mouse-Fibroblast datasets, we performed independent RSEM simulations: we first performed isoform quantification for each cell, followed by read simulation based on RSEM’s inferred expression estimates from latent variables. This two-step process ensured that each simulated cell reflected the expression characteristics of its corresponding real cell. The resulting simulated datasets, HEK293T-Sim with 192 human cells and Fibroblast-Sim with 369 mouse cells, were evaluated across the same range of cell scales as their real counterparts.

#### Ground-Truth for Evaluation

For real datasets, since the true expressed transcripts for each cell are unknown, we used the reference annotation (Ensembl GRCh38.107 for human and Ensembl GRCm39.110 for mouse) as the ground-truth. We acknowledge that using the entire transcriptome as reference may overestimate cell-specific assembly accuracy, as transcripts not expressed in a cell may be considered correct if annotated in the reference. Nevertheless, this approach still provides a fair comparison of the tools’ relative accuracy. For simulated datasets, each cell has its own, distinct expressed transcripts, serving as ground-truth for rigorous evaluation of assembly methods, in particular their cell-specific assembly accuracy.

#### Evaluation Metrics

As a common practice, we defined an assembled multi-exon transcript as “matching” if its intron-chain exactly matched that of a transcript in the reference ground-truth. We focused on multiexon transcripts as they are biologically more interesting while ensuring fair comparison with TransMeta and Aletsch, which only assemble multi-exon transcripts. With these, we use two metrics: the number of matching transcripts, which is proportional to sensitivity, and precision, defined as the ratio of matching transcripts to total assembled transcripts. The tool GffCompare [16] was used to calculate these two metrics. In cases where two methods demonstrate different trade-offs between precision and sensitivity (i.e., one method gives higher sensitivity but lower precision), we compare their precision at the same level of sensitivity, known as adjusted precision [19,32]. The adjusted precision of method *X* w.r.t. another method *Y* is calculated by gradually filtering out the lower-scoring transcripts from *X* until it matches the sensitivity of *Y*. This is equivalent to locating the point on the precision-recall curve of *X* that has the same sensitivity as *Y*.

### 3.2 Comparison on Real Single-cell RNA-seq Datasets

Figs. 2 and 3 compare assembly accuracy of the six methods. Beaver demonstrated superior performance, achieving the highest precision and recall in both datasets. On the HEK293T dataset, Beaver marginally surpassed TransMeta, the second-best method, in precision (83% vs. 82%) while exhibiting substantially higher sensitivity (68.4% more transcripts on median); on the Mouse-Fibroblast dataset, Beaver outperformed TransMeta by a large margin in both precision and recall.

**Fig. 2:**
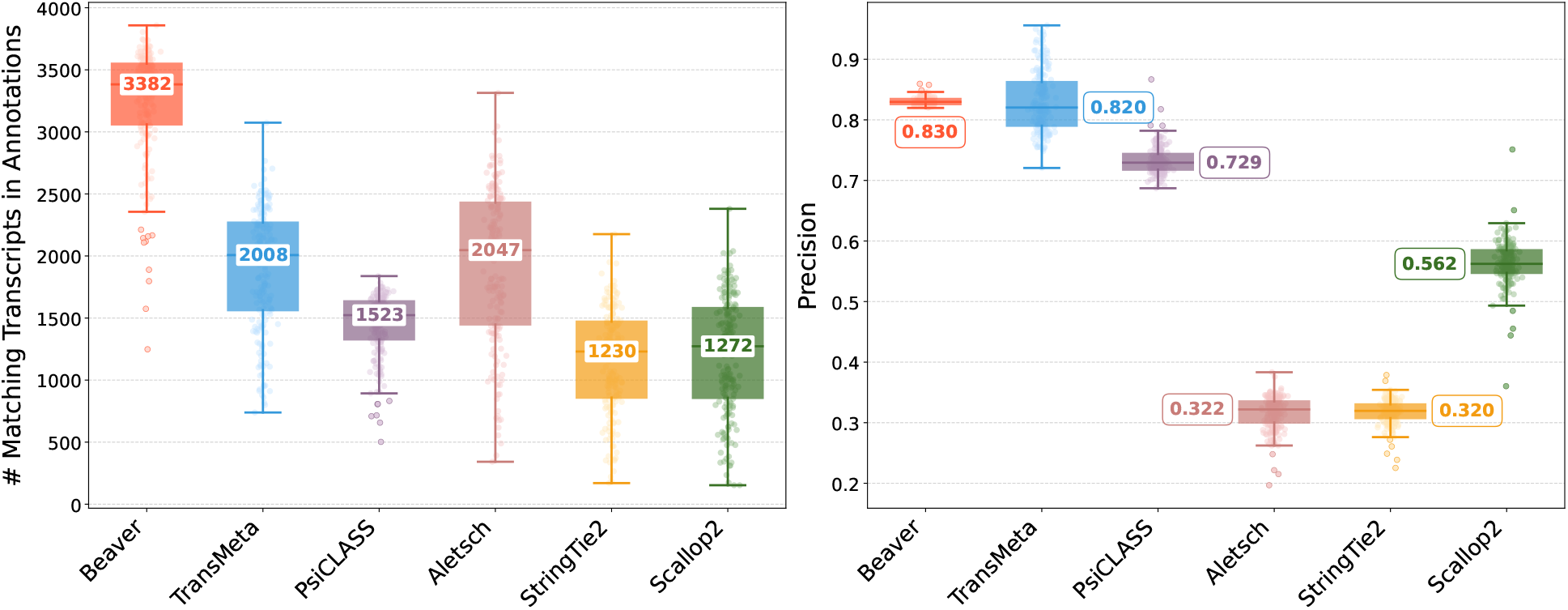
Comparison of assembly performance on the HEK293T dataset. Left: number of matching transcripts; right: precision. Median values are annotated.

**Fig. 3:**
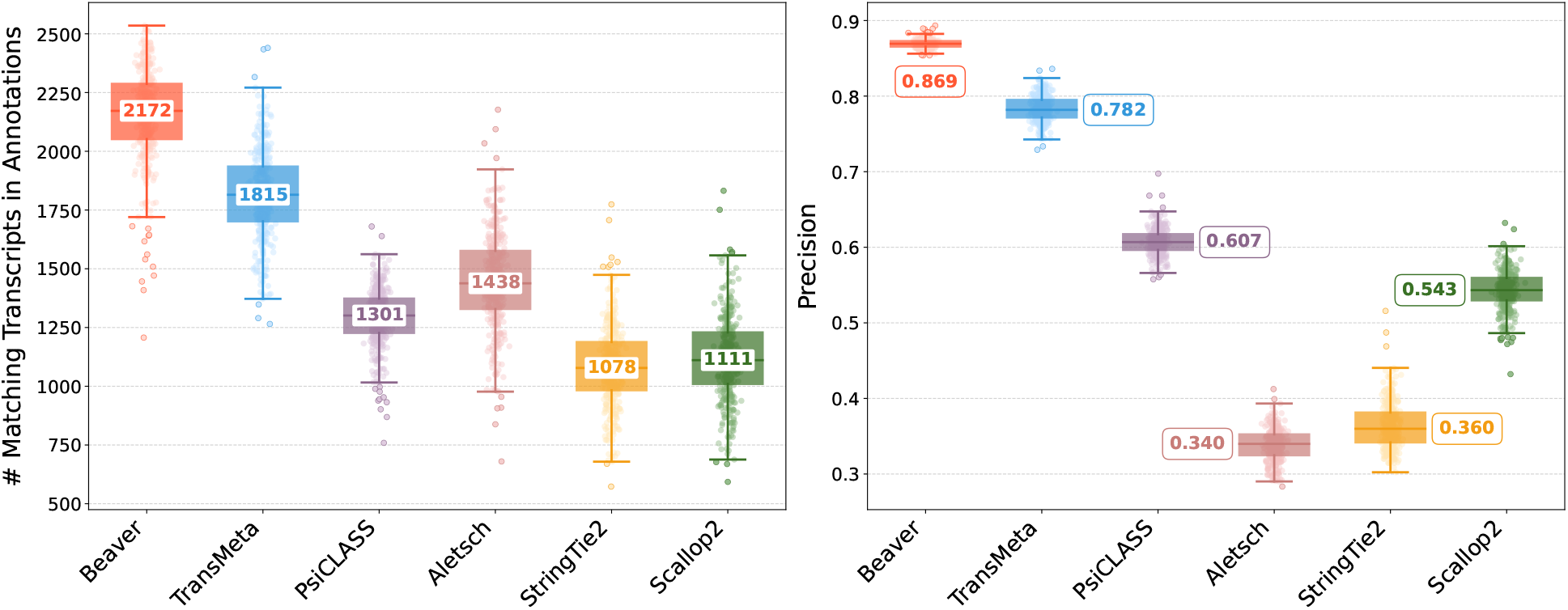
Comparison of assembly performance on the Mouse-Fibroblast dataset. Left: number of matching transcripts; right: precision. Median values are annotated.

We further evaluated the adjusted precision between Beaver and each of the other five methods. Fig. 4 and Fig. 5 display these comparisons at the single-cell level (comparisons across all scales are provided in Supplementary Figure 1–10). We observe that Beaver’s points consistently lie to the right of competing methods, indicating remarkable precision across all cells at equivalent sensitivity levels. Table 1 summarizes the mean adjusted precision across all cells. The improvement over Aletsch, whose individual assemblies served as Beaver’s input, was particularly noteworthy: 64.6% for the HEK293T dataset and 62.2% for the Mouse-Fibroblast datasets. These improvements clearly validate the effectiveness of Beaver’s innovative techniques.

**Table 1:** Comparison of adjusted precision (%) averaged over all cells in each real dataset. Abbreviations: TM = TransMeta; PC = PsiCLASS; AT = Aletsch; ST2 = StringTie2; SC2 = Scallop2; BV = Beaver.

| Dataset | TM vs. BV |  |  | PC vs. BV |  |  | AT vs. BV |  |  | ST2 vs. BV |  |  | SC2 vs. BV |  |  |
| --- | --- | --- | --- | --- | --- | --- | --- | --- | --- | --- | --- | --- | --- | --- | --- |
| | TM | BV | $\Delta\%$ | PC | BV | $\Delta\%$ | AT | BV | $\Delta\%$ | ST2 | BV | $\Delta\%$ | SC2 | BV | $\Delta\%$ |
| HEK293T | 83.0 | 96.5 | 13.5 | 73.3 | 98.2 | 24.9 | 31.6 | 96.2 | 64.6 | 31.8 | 98.8 | 67.0 | 56.4 | 98.7 | 42.3 |
| Fibroblast | 78.3 | 92.6 | 14.3 | 60.8 | 97.1 | 36.3 | 33.8 | 96.0 | 62.2 | 36.4 | 98.0 | 61.7 | 54.3 | 97.9 | 43.6 |

**Fig. 4:**
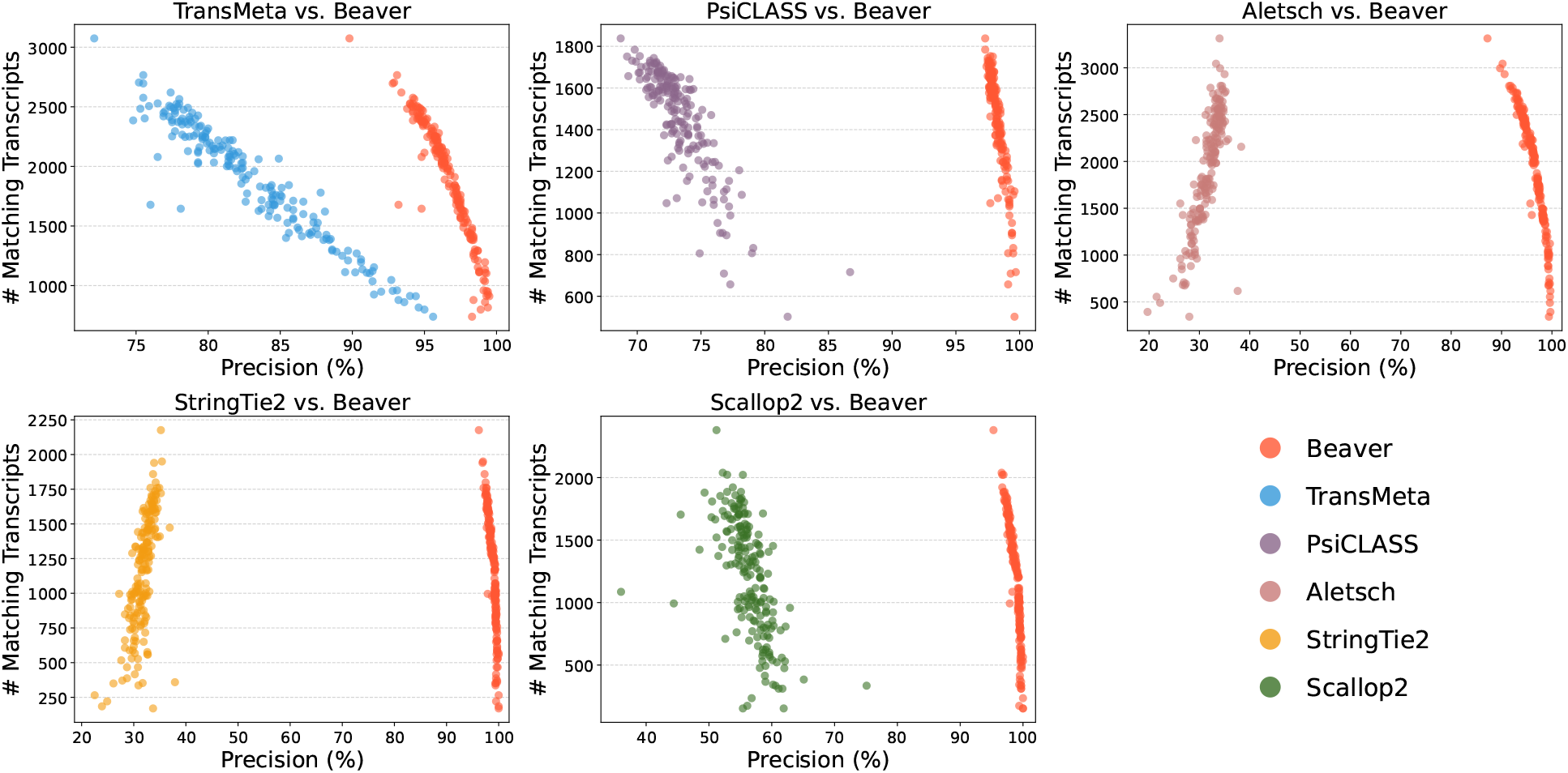
Pairwise comparison of adjusted precision across individual HEK293T cells (*n* = 192).

**Fig. 5:**
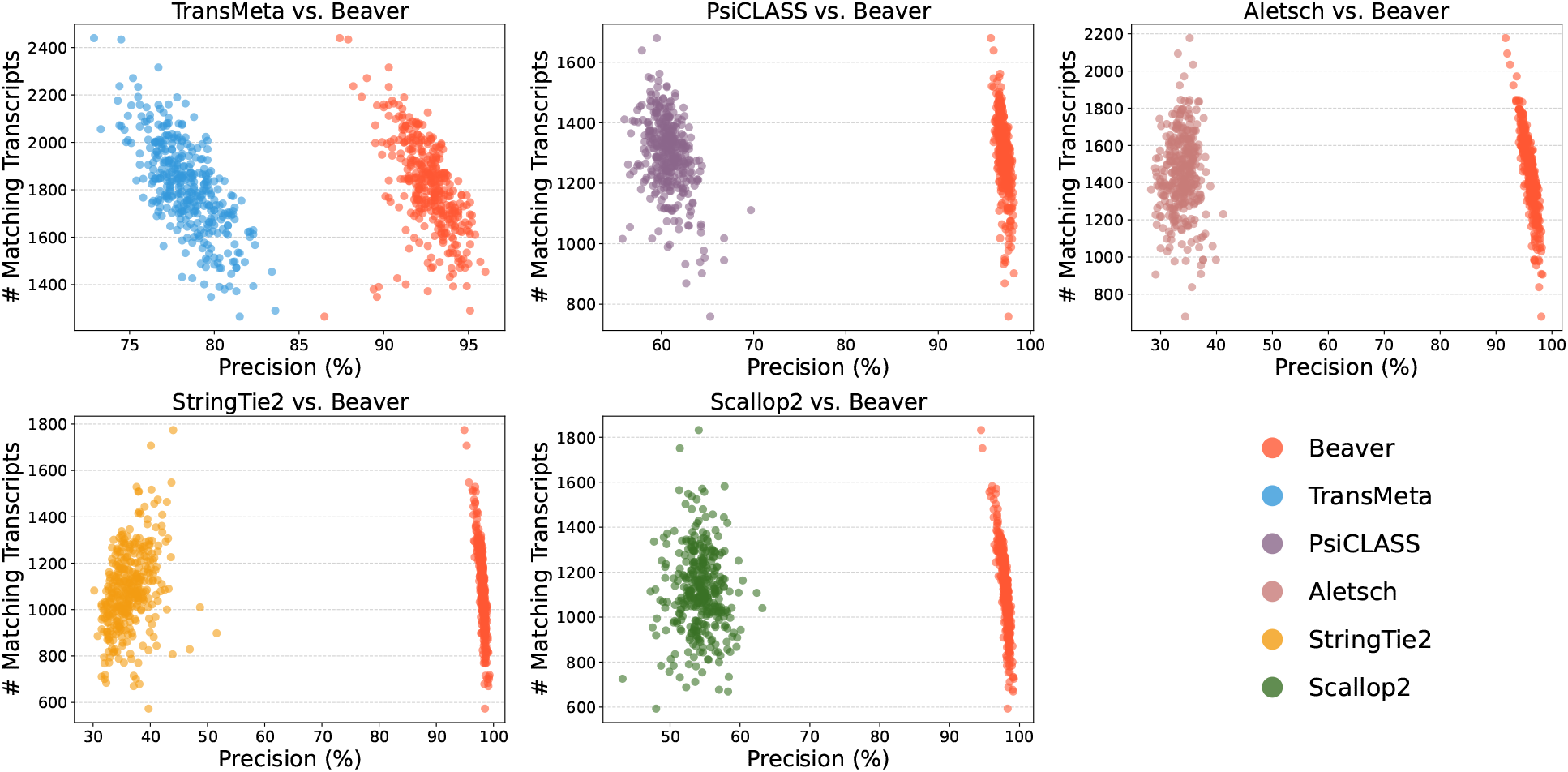
Pairwise comparison of adjusted precision across individual Mouse-Fibroblast cells (*n* = 369).

It is important to note that real datasets lack true cell-specific expressed transcripts—the ground-truth (i.e., reference annotation) is the same for all cells. Although this approach may not fully capture cell-specific expression patterns, it demonstrates Beaver’s enhanced capability to identify reliable transcripts previously verified in an annotation. We use simulations to evaluate cell-specific assembly (Section 3.3).

### 3.3 Comparison on Simulated Single-cell RNA-seq Datasets

Figs. 6 and 7 compare the assembly accuracy of different methods on the two simulated datasets. Again, Beaver achieved the marked sensitivity while maintaining highest precision. Table 2 and Figs. 8, 9 present the comparison of adjusted precision, showing substantial improvements by Beaver over all other methods (detailed comparisons across all cell scales are available in Supplementary 11–20). Given that the simulated datasets are cell-specific, with true expressed transcripts varying across cells, these results provide strong evidence of Beaver’s superiority over other methods in generating accurate assemblies at single-cell resolution.

**Table 2:** Comparison of adjusted precision (%) averaged over all cells in the simulated dataset. Abbreviations: TM = TransMeta; PC = PsiCLASS; AT = Aletsch; ST2 = StringTie2; SC2 = Scallop2; BV = Beaver.

| Dataset | TM vs. BV |  |  | PC vs. BV |  |  | AT vs. BV |  |  | ST2 vs. BV |  |  | SC2 vs. BV |  |  |
| --- | --- | --- | --- | --- | --- | --- | --- | --- | --- | --- | --- | --- | --- | --- | --- |
| | TM | BV | $\Delta\%$ | PC | BV | $\Delta\%$ | AT | BV | $\Delta\%$ | ST2 | BV | $\Delta\%$ | SC2 | BV | $\Delta\%$ |
| HEK293T-Sim | 54.4 | 91.0 | 36.6 | 72.4 | 88.3 | 15.9 | 49.4 | 81.4 | 32.0 | 58.5 | 88.7 | 30.2 | 72.1 | 85.6 | 13.5 |
| Fibroblast-Sim | 62.1 | 93.1 | 31.0 | 83.3 | 93.1 | 9.8 | 58.5 | 90.6 | 32.1 | 68.9 | 93.2 | 24.3 | 82.1 | 92.2 | 10.1 |

**Fig. 6:**
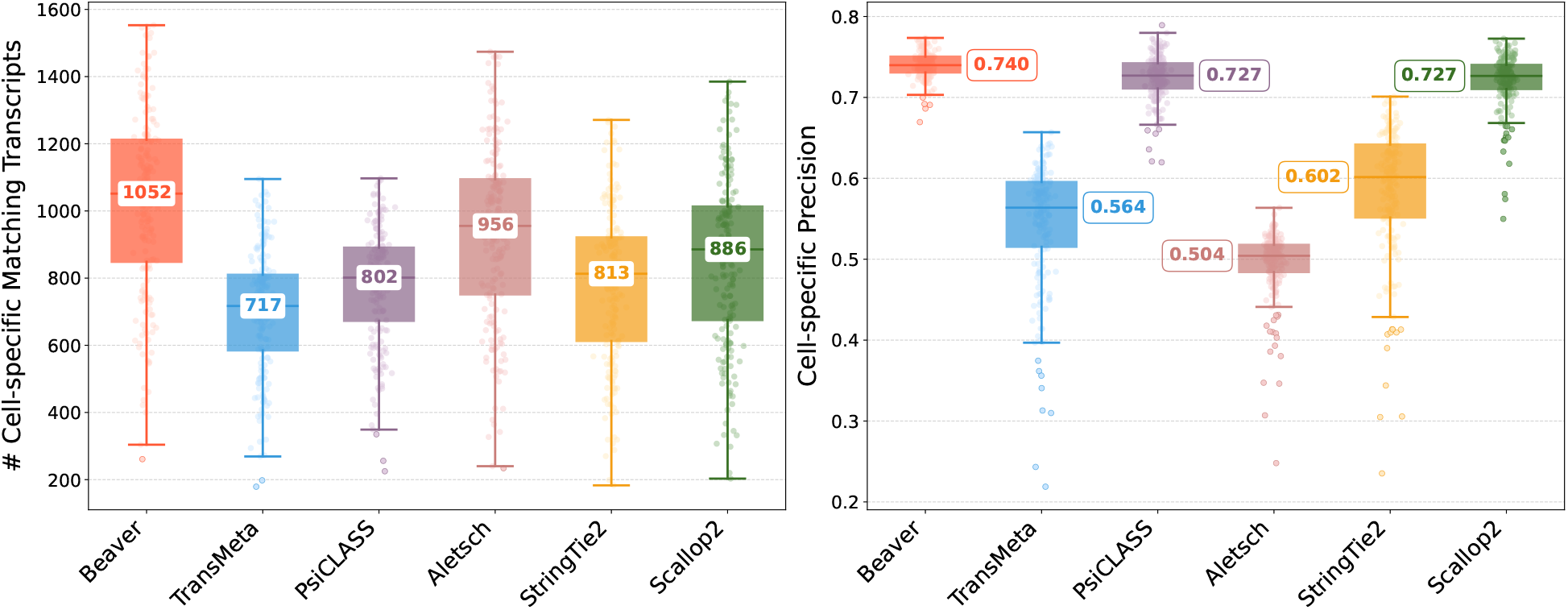
Comparison of assembly performance on the HEK293T-Sim dataset. Left: number of matching transcripts; right: precision. Median values are annotated.

**Fig. 7:**
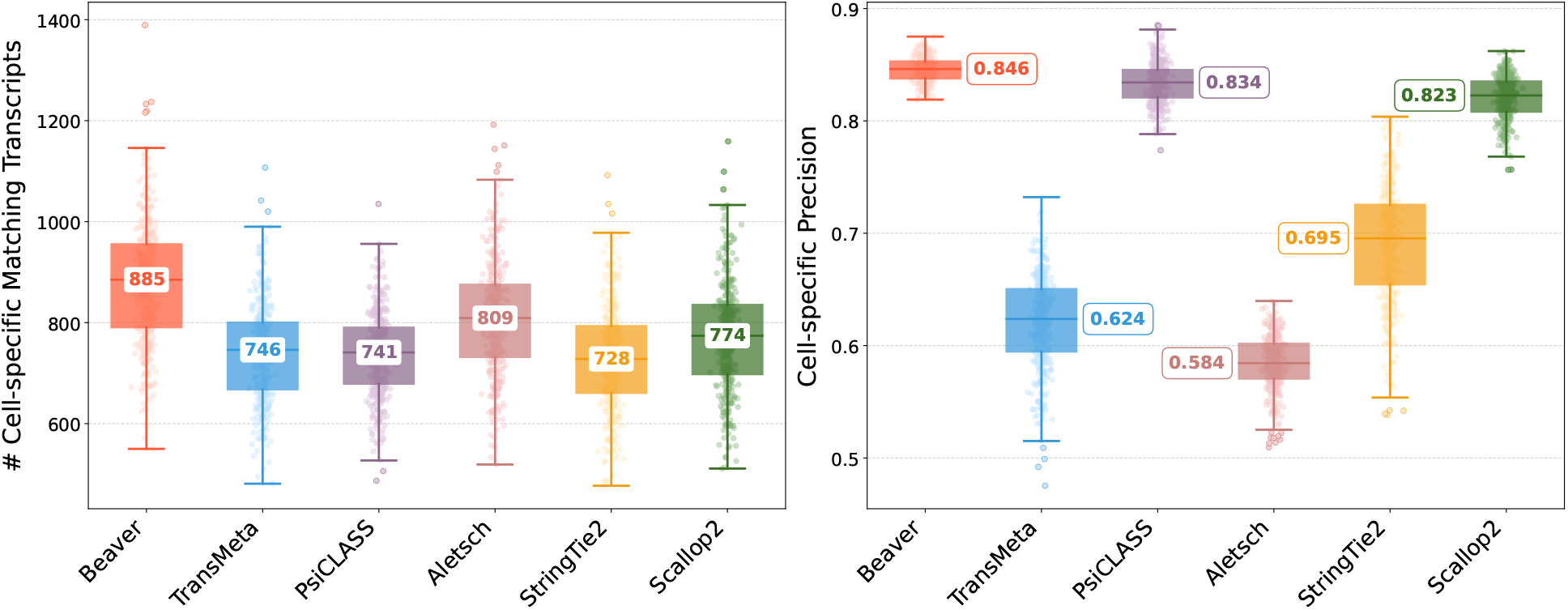
Comparison of assembly performance on the Fibroblast-Sim dataset. Left: number of matching transcripts; right: precision. Median values are annotated.

**Fig. 8:**
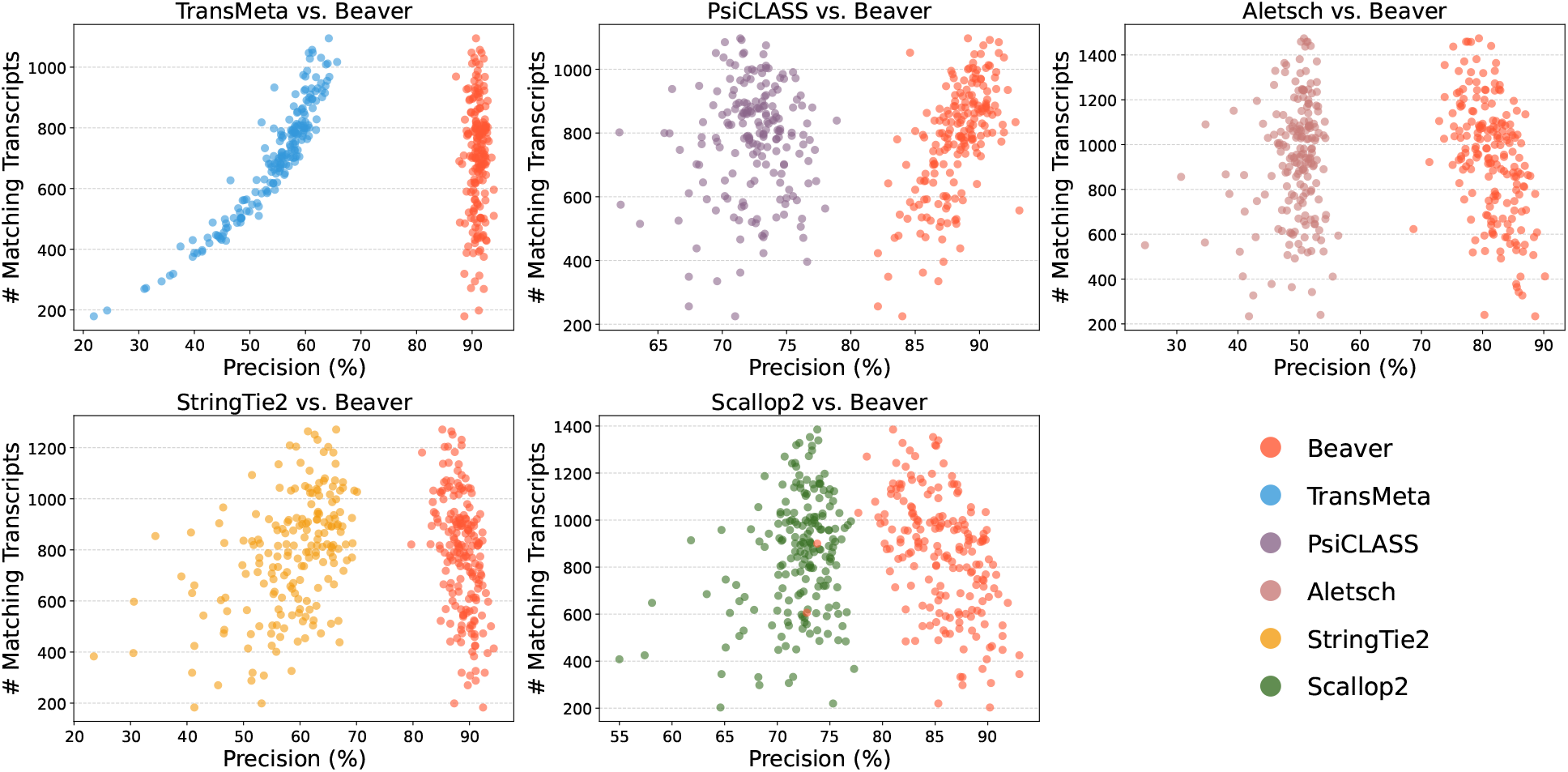
Pairwise comparison of adjusted precision across individual HEK293T-Sim cells (*n* = 192).

**Fig. 9:**
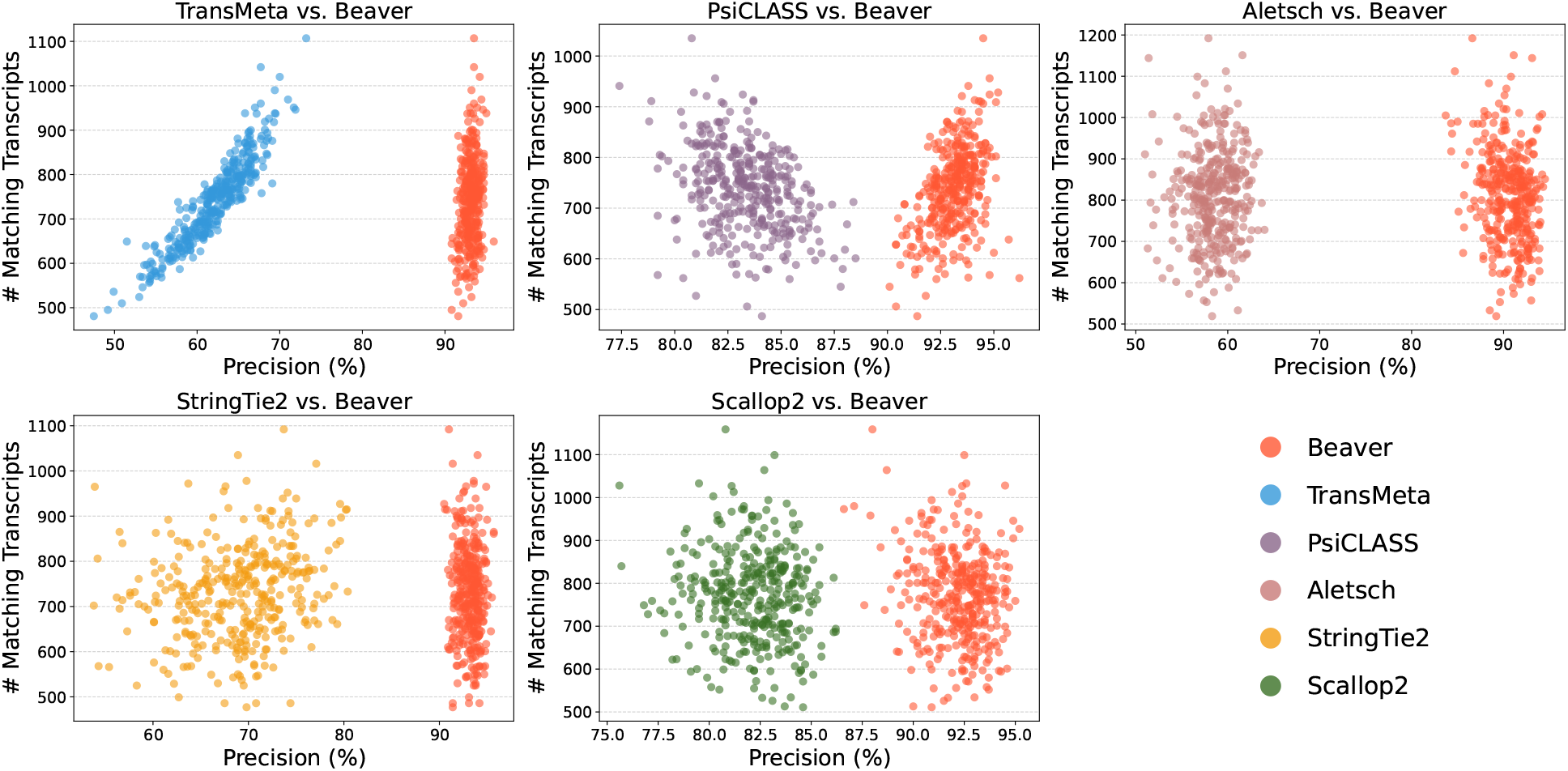
Pairwise comparison of adjusted precision across individual Fibroblast-Sim cells (*n* = 369).

We observed clear divergence in methods’ performance between real and simulated datasets. For example, TransMeta, a meta-assembler, ranked second on real datasets but showed relatively poor performance on simulated datasets, particularly for cells with lower transcript expression. Conversely, Scallop2, a single-sample assembler, did not stand out in precision on real datasets yet achieved nearly the highest precision on simulated datasets. To investigate this discrepancy using the known cell-specific ground truth, we analyzed two categories of matching transcripts: “cell-specific” matches (those matching individual cell ground truth) and “general” matches (those matching any transcript in the collective ground truth across all cells). Fig. 10 presents the mean number of predicted transcripts across all cells in the simulated dataset.

**Fig. 10:**
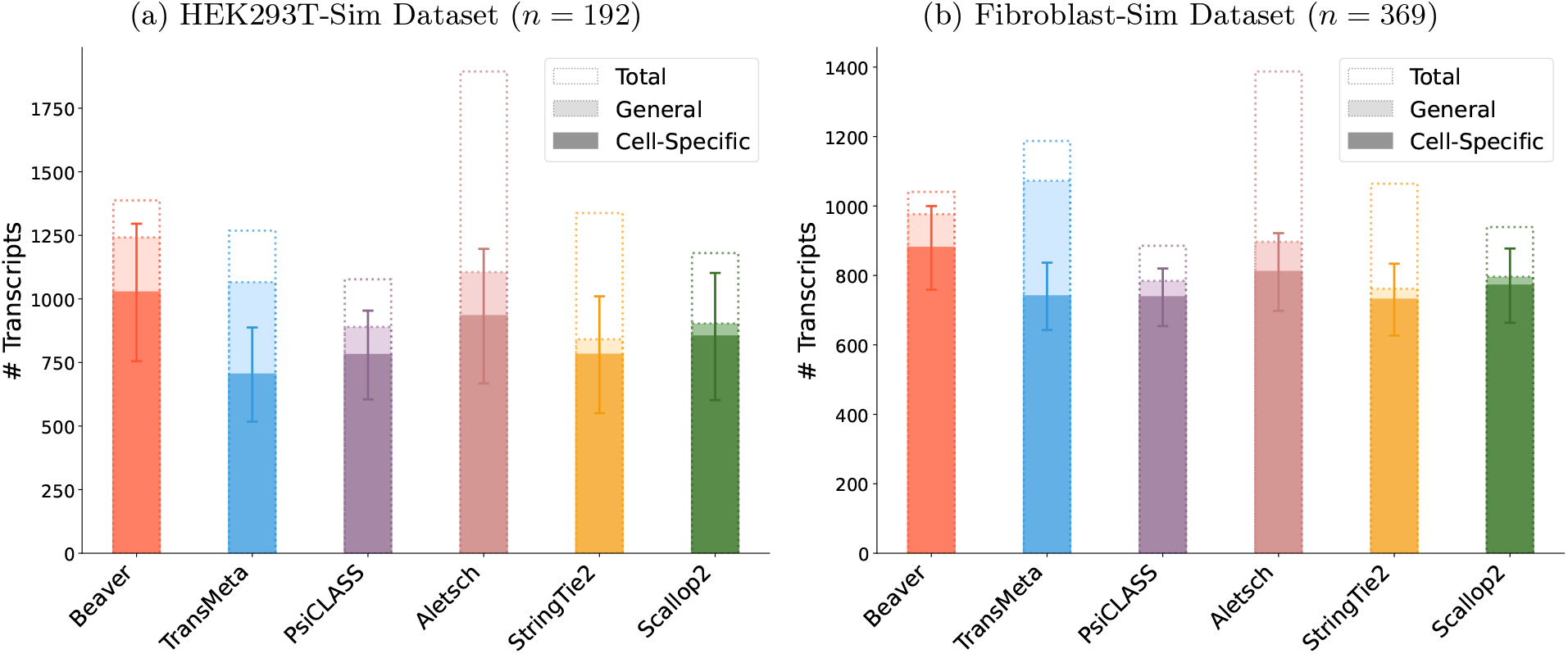
Comparison of “General” versus “Cell-Specific” matching counts across assemblers in simulated datasets. Single-sample assemblers show minimal divergence between matching types. Beaver significantly reduces false positives from Aletsch’s individual assemblies (indicated by dotted boundaries).

Single-sample assemblers (e.g., StringTie2 and Scallop2), which do not access data from other cells, showed minimal divergence between general and cell-specific matching. In contrast, meta-assemblers, designed to generate accurate meta-assemblies for all cells, often struggled with precise transcript assignment to individual cells. TransMeta, for instance, uses a strategy where assembled transcripts are assigned to cells if they cover half of a transcript’s junctions, leading to overestimation of transcript presence in individual cells and creating substantial gaps between general and cell-specific matching accuracy. Beaver adopts a distinct approach, achieving a superior balance by first aggressively assigning reliable transcripts to individual assemblies, followed by using comprehensive cell-specific features for scoring. This strategy fully leverages shared information across cells while achieving significantly improved cell-specific accuracy.

## 4 Conclusion and Discussion

We introduce Beaver, a new transcript assembler for scRNA-seq data that substantially improves accuracy at single-cell resolution. Beaver’s methodological innovations include a graph-based data structure and dynamic programming algorithm that effectively reconstructs candidate full-length transcripts from incomplete and fragmented individual assemblies. This is followed by precise cell-specific scoring using a two-stage machine learning model trained on 51 custom-designed features. Our experiments demonstrate that Beaver successfully addresses the challenge of missing junctions in scRNA-seq data, accurately producing full-length assemblies that capture cellular heterogeneity.

Beaver’s approach to transcript reconstruction, which aggregates individual cell assemblies to generate fulllength transcripts, conceptually parallels the transcript recovery methods in long-read transcript assembly. Beaver takes advantage of the high-accuracy in the splicing position offered by short-reads scRNA-data, meanwhile addresses the major challenge posed by coverage gaps. While long-read sequencing technologies, such as PacBio Iso-Seq and Oxford Nanopore, offer the capability to sequence full-length transcripts [15], their widespread adoption has been limited in single cells by relatively low throughput, high error rates and high costs [2,21,8]. Currently, short-read sequencing remains dominant for single-cell transcriptomics. The development of hybrid approaches that combine short-read and long-read data [26] represents a promising direction for isoform detection and quantification at single-cell resolution. Future iterations of Beaver could incorporate long-read guidance for transcript selection to minimize inappropriate junction combinations and better preserve cell-specific splicing patterns.

The effective training of Beaver on real scRNA-seq data is limited by the lack of high-quality datasets with known expressed transcripts. In this study, we used reference annotations as ground truth for two real RNA-seq datasets, but these are not cell-specific. The high accuracy and significant improvements observed on simulated data (where cell-specific transcripts are known) demonstrate that an accurate model *can* be trained when such data is available. One of our future work will focus on collecting real scRNA-seq data with curated, cell-specific transcripts, to develop even more accurate models.

We anticipate that Beaver will become a widely used tool for scRNA-seq analysis. One direct application is the identification of novel isoforms expressed in specific cell or cell type. While tools like alevin-fry [5] and kallisto|bustools [24] have made significant advances in gene-level quantification for single cells, transcript-level abundance estimation remains challenging [6]. By providing a more accurate and cell-specific transcriptome references, Beaver can potentially enhance both the accuracy and computational efficiency of existing quantification methods.

## Acknowledgments

This work is supported by the US National Science Foundation (DBI-2145171 to M.S.) and the US National Institutes of Health (R01HG011065 to M.S.).

