## Supplementary Materials for "Transcriptome Assembly at Single-Cell Resolution with Beaver"

### Supplementary Materials for
### "Transcriptome Assembly at Single-Cell Resolution with Beaver"

Qian Shi[1], Qimin Zhang[1], and Mingfu Shao[1,2,*]

[1]Department of Computer Science and Engineering, School of Electronic Engineering and Computer Science, The Pennsylvania State University

[2]Huck Institutes of the Life Sciences, The Pennsylvania State University

October 25, 2024

#### List of Supplementary Notes

1 PseudoCode for Candidate Transcript Generation . . . 3
2 Features Description . . . 4

#### List of Supplementary Figures

1 Beaver vs. TransMeta on HEK293T dataset. . . . 7
2 Beaver vs. PsiCLASS on HEK293T dataset. . . . 7
3 Beaver vs. Aletsch on HEK293T dataset. . . . 8
4 Beaver vs. StringTie2 on HEK293T dataset. . . . 8
5 Beaver vs. Scallop2 on HEK293T dataset. . . . 9
6 Beaver vs. TransMeta on Mouse-Fibroblast dataset. . . . 9
7 Beaver vs. PsiCLASS on Mouse-Fibroblast dataset. . . . 10
8 Beaver vs. Aletsch on Mouse-Fibroblast dataset. . . . 10
9 Beaver vs. StringTie2 on Mouse-Fibroblast dataset. . . . 11
10 Beaver vs. Scallop2 on Mouse-Fibroblast dataset. . . . 11
11 Beaver vs. TransMeta on HEK293T-Sim dataset. . . . 12

12 Beaver vs. PsiCLASS on HEK293T-Sim dataset. . . . . . . . . . . . . . . . . . . . . . . . . . . . . 12
13 Beaver vs. Aletsch on HEK293T-Sim dataset. . . . . . . . . . . . . . . . . . . . . . . . . . . . . . 13
14 Beaver vs. StringTie2 on HEK293T-Sim dataset. . . . . . . . . . . . . . . . . . . . . . . . . . . . . 13
15 Beaver vs. Scallop2 on HEK293T-Sim dataset. . . . . . . . . . . . . . . . . . . . . . . . . . . . . . 14
16 Beaver vs. TransMeta on Mouse-Fibroblast-Sim dataset. . . . . . . . . . . . . . . . . . . . . . . . . 14
17 Beaver vs. PsiCLASS on Mouse-Fibroblast-Sim dataset. . . . . . . . . . . . . . . . . . . . . . . . . 15
18 Beaver vs. Aletsch on Mouse-Fibroblast-Sim dataset. . . . . . . . . . . . . . . . . . . . . . . . . . 15
19 Beaver vs. StringTie2 on Mouse-Fibroblast-Sim dataset. . . . . . . . . . . . . . . . . . . . . . . . 16
20 Beaver vs. Scallop2 on Mouse-Fibroblast-Sim dataset. . . . . . . . . . . . . . . . . . . . . . . . . 16

### Supplementary Note 1: PseudoCode for Transcript Generation

**Algorithm 1** Dynamic Programming for Candidate Transcript Generation

**Input:** Connected component $G_c = (V_c, E_c)$ ▷ Connected components n transcript fragment graph
**Output:** Set of optimal paths $P_c$ ▷ Each path represents a candidate transcript

```
function FindOptimalPaths($G_c$)
    $(v_1, v_2, \ldots, v_{|V_c|}) \leftarrow$ TopologicalSort($G_c$)    ▷ $O(|V_c| + |E_c|)$
    paths[$v$] $\leftarrow$ new MinHeap() for all $v \in V_c$    ▷ Initialize Min-Heaps
    for $j \leftarrow 1$ to $|V_c|$ do
        paths[$v_i$].push{(MergingScore($v_i$), $v_i$}    ▷ Initialize single-vertex path
        for $(v_i, v_j) \in E_c$ do    ▷ Check incoming edges to $v_j$
            for each path $p$ in paths[$v_i$].getAll() do    ▷ $O(p_n)$
                new_path $\leftarrow p \cup \{v_j\}$    ▷ Extend path with new vertex
                score $\leftarrow$ MergingScore(new_path)
                if paths[$v_j$].size() $< p_n$ then    ▷ Limit $p_n$ paths stored per node
                    paths[$v_j$].push(score, new_path)
                else if score $>$ paths[$v_j$].top().score then    ▷ Heap is full
                    paths[$v_j$].pop()    ▷ Remove lowest scoring path in the Min-Heap
                    paths[$v_j$].push(score, new_path)
                end if
            end for
        end for
    end for
    return $P_c \leftarrow$ TopK($\bigcup_{v \in V_c} \{p \mid (\text{score}, p) \in \text{paths}[v].\text{getAll}()\}, p_c$)    ▷ Limit $p_c$ paths stored per graph
end function
function MergingScore(path $p$)
    $b \leftarrow \min_{j \in J(p)}$ JunctionWeight($j, p$)    ▷ Bottleneck weight
    $n \leftarrow |J(p)|$    ▷ Number of junctions
    return $b \cdot n$
end function
function JunctionWeight(junction $j$, path $p$)
    $X_j \leftarrow \{t \mid t$ contains $j$ and is compatible with $p\}$    ▷ Find compatible transcripts
    return $\sum_{t \in X_j}$ coverage($t$)    ▷ Sum coverage of supporting transcripts
end function
```

#### Supplementary Note 2: Features Description

This note describes the feature sets used to train Beaver-General and Beaver-Specific. We developed two feature sets: (1) general features (n=30) that assess overall transcript reliability across the dataset, and (2) cell-specific features (n=21) that evaluate expression likelihood in individual cells. These features are computed for every unique pair of candidate path $p$ and cell $c$ in the transcript fragment graphs, with general features being replicated across all cells for the same $p$.

(Shown in *Italic* are the headers of the Beaver's feature table.)

**General Features ($n = 30$)**

- *sample_size*: Total number of input cells/samples.
- *bottleneck_coverage*: Bottleneck junction-score $BJ(p)$ from the dynamic programming solution.
- *highest_coverage*: Maximum coverage value observed among raw transcript fragments $t \sim p$.
- *num_junctions*: Total number of junctions in path $p$, i.e. $NJ(p)$.
- Compatible junction coverage:

  (*min_comp_cov*, *max_comp_cov*, *median_comp_cov*, *mean_comp_cov*, *std_comp_cov*)

  Statistics of junction-scores defined in Section 2.3, which is the sum of scores of all transcript fragments that contain junction $j$ and are compatible with $p$, i.e., $J(j, p) := \sum_{t:j \in t \text{ and } t \sim p} score(t)$.
- Junction coverage statistics:

  (*min_junc_cov*, *max_junc_cov*, *median_junc_cov*, *mean_junc_cov*, *std_junc_cov*)

  Statistics of junction-scores regardless of compatibility, i.e., $J'(j, p) := \sum_{t:j \in t} score(t)$.
- *extendable_score*: Cumulative junction scores for potential left/right path extensions.
- *cells_support*: Number of cells supporting path $p$.
- *ratio_cells_support*: Proportion of total cells supporting path $p$.
- *ratio_cells_fl_over_sample*: Proportion of cells supporting all junctions in $p$.

- *ratio_cells_fl_over_cells_support*: Proportion of full-length support among supporting cells.

- Cell support statistics:

  (*min_cell_junc*, *max_cell_junc*, *median_cell_junc*, *mean_cell_junc*, *std_cell_junc*)

  Statistics describing the number of cells supporting individual junctions $j \in p$.

- *num_fragments*: Total number of fragments compatible with $p$.

- Fragment Connecting:

  (*min_frag_cov*, *max_frag_cov*, *median_frag_cov*, *mean_frag_cov*, *std_frag_cov*)

  Statistics describing fragment coverage along path $p$.

**Cell-Specific Features ($n = 21$)**

- *input_genes*: Estimated number of input gene loci in cell $c$.

- *input_transcripts*: Number of input transcripts in cell $c$.

- *output_transcripts*: Number of output transcripts in cell $c$.

- *cell_supporting_junctions*: Number of junctions $j \in p$ supported by $c$.

- *cell_ratio_supporting_junctions*: Proportion of junctions $j \in p$ is supported by $c$.

- Cell-specific compatible junction coverage:

  (*cell_min_comp_cov*, *cell_max_comp_cov*, *cell_median_comp_cov*, *cell_mean_comp_cov*, *cell_std_comp_cov*)

  Statistics of cell-specific junction-scores, i.e., $J(c, p, j) := \sum_{t:j \in t \text{ and } t \sim p \text{ and } t \in c} score(t)$.

- *cell_nonzero_comp_cov*: Number of junction $j \in p$ such that $J(c, p, j) \neq 0$.

- *cell_max_streak_comp_cov*: Maximum number of continuous non-zero $J(c, p, j)$ for all junctions $j \in p$.

- *cell_min_streak_comp_cov*:Minimum number of continuous non-zero $J(c, p, j)$ for all junctions $j \in p$.

- Cell-specific Junction coverage:

  (*cell_min_junc_cov*, *cell_max_junc_cov*, *cell_median_junc_cov*, *cell_mean_junc_cov*, *cell_std_junc_cov*)

Statistics of cell-specific junction-scores regardless of compatibility, i.e., $J'(c,p,j) := \sum_{t:j \in t \text{ and } t \in c} score(t)$.

- *cell_nonzero_junc_cov*: Number of junction $j \in p$ such that $J'(c,p,j) \neq 0$.
- *cell_max_streak_junc_cov*: Maximum number of continuous non-zero $J'(c,p,j)$ for all junctions $j \in p$.
- *cell_min_streak_junc_cov*: Minimum number of continuous non-zero $J'(c,p,j)$ for all junctions $j \in p$.

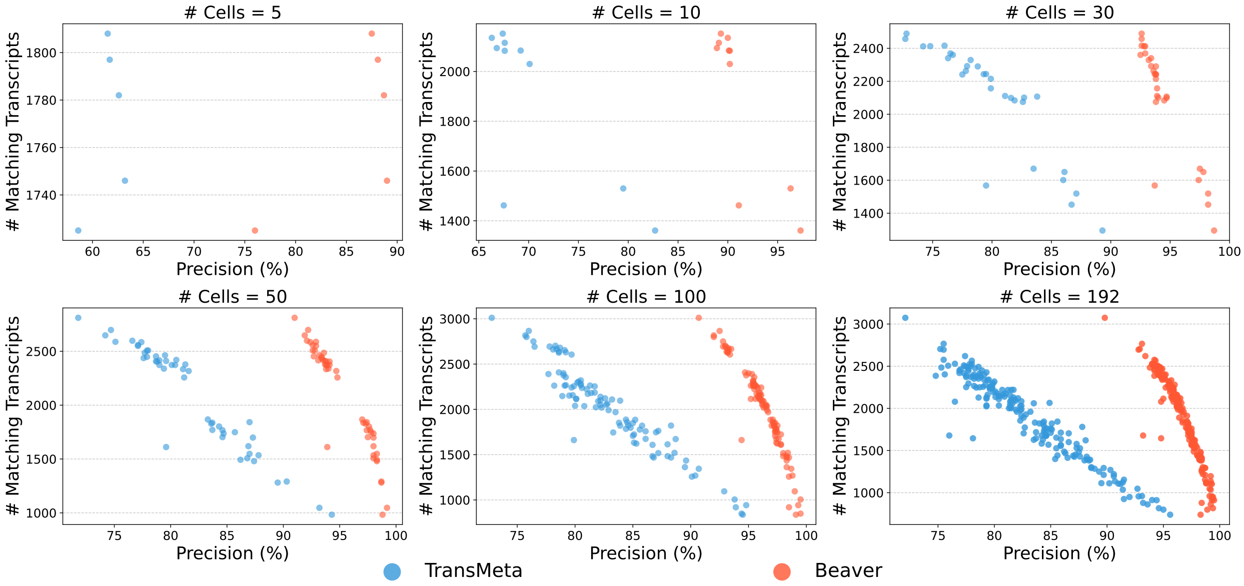


**Supplementary Figure 1**: Comparison of adjusted precision between Beaver and TransMeta across varying cell population sizes ($n = 5, 10, 30, 50, 100, 192$) in the HEK293T dataset. Each data point represents an individual cell.

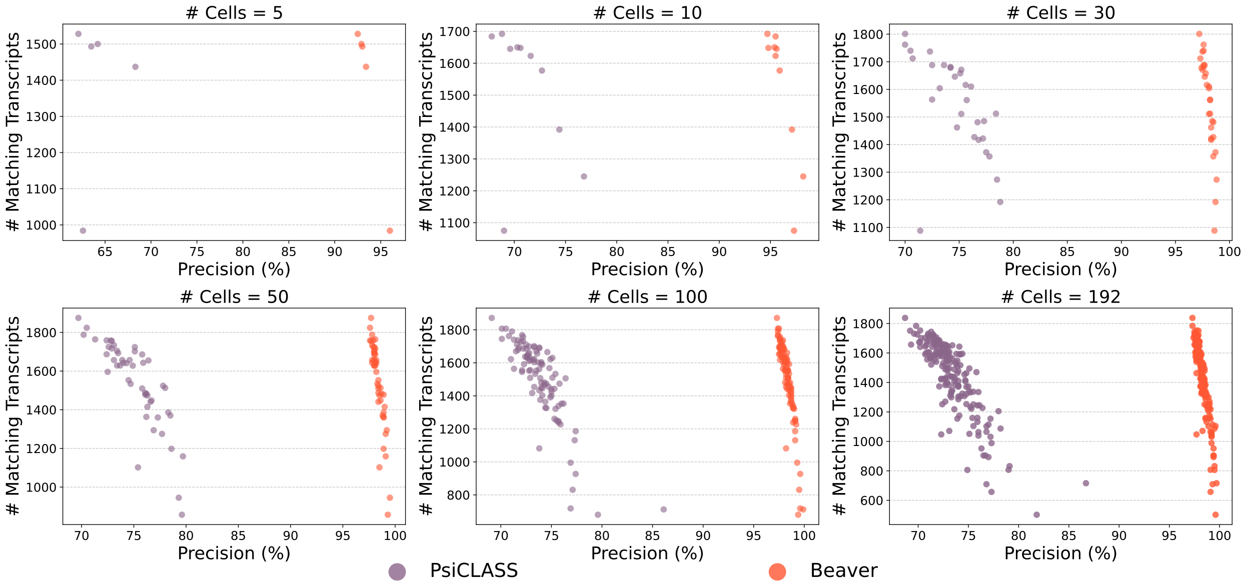


**Supplementary Figure 2**: Comparison of adjusted precision between Beaver and PsiCLASS across varying cell population sizes ($n = 5, 10, 30, 50, 100, 192$) in the HEK293T dataset. Each data point represents an individual cell.

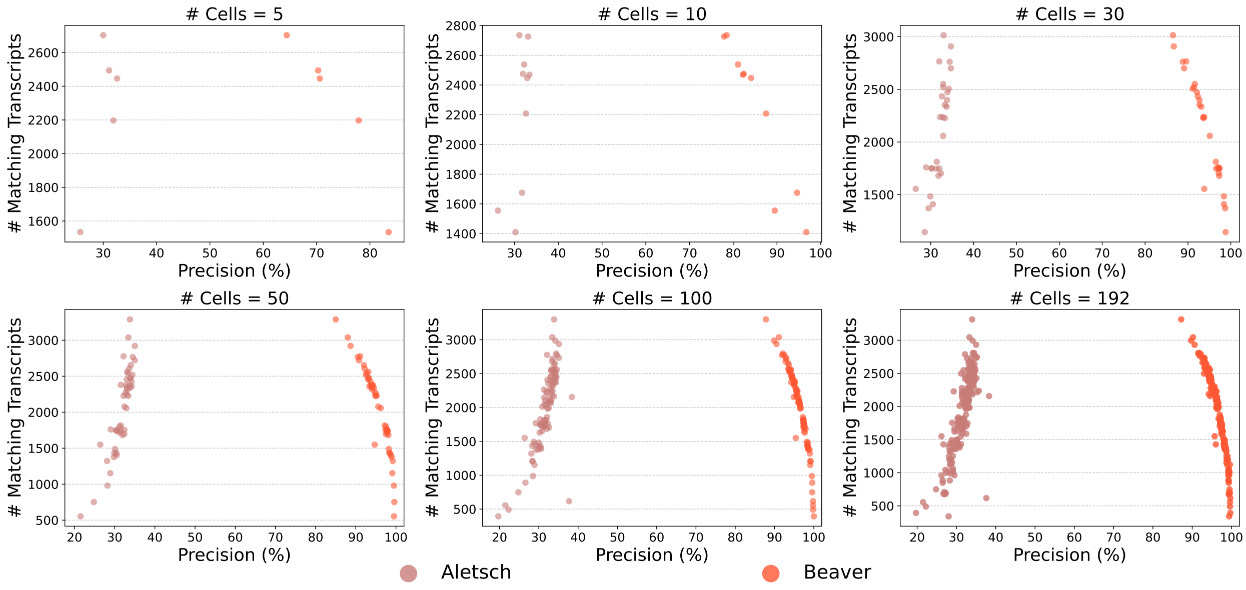


**Supplementary Figure 3**: Comparison of adjusted precision between Beaver and Aletsch across varying cell population sizes ($n = 5, 10, 30, 50, 100, 192$) in the HEK293T dataset. Each data point represents an individual cell.

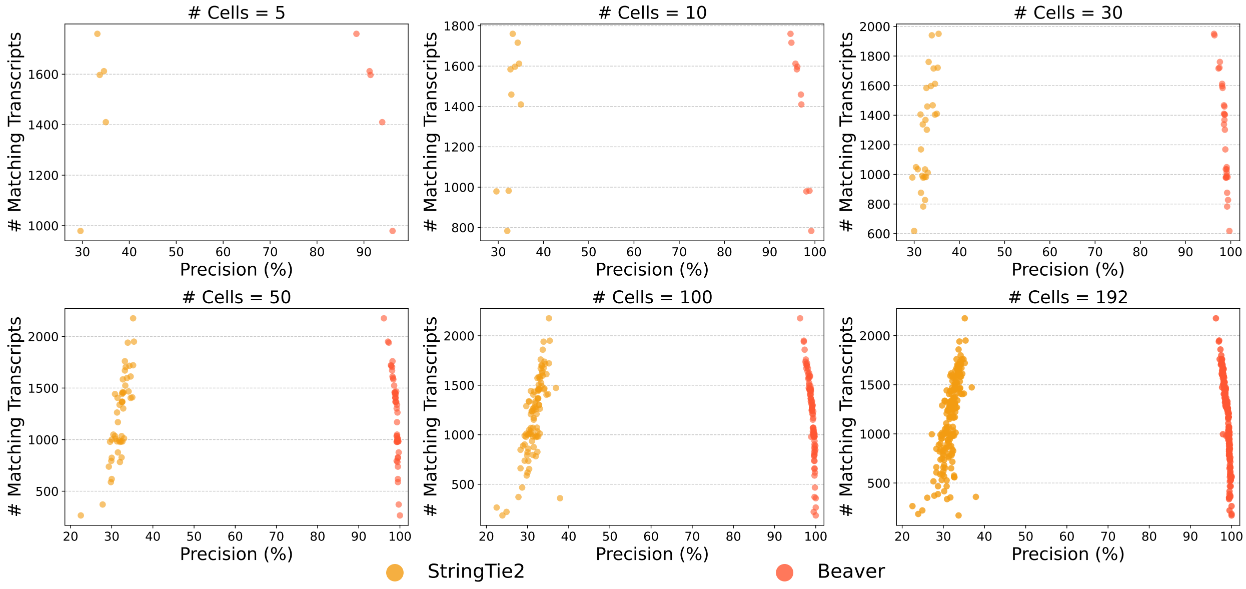


**Supplementary Figure 4**: Comparison of adjusted precision between Beaver and StringTie2 across varying cell population sizes ($n = 5, 10, 30, 50, 100, 192$) in the HEK293T dataset. Each data point represents an individual cell.

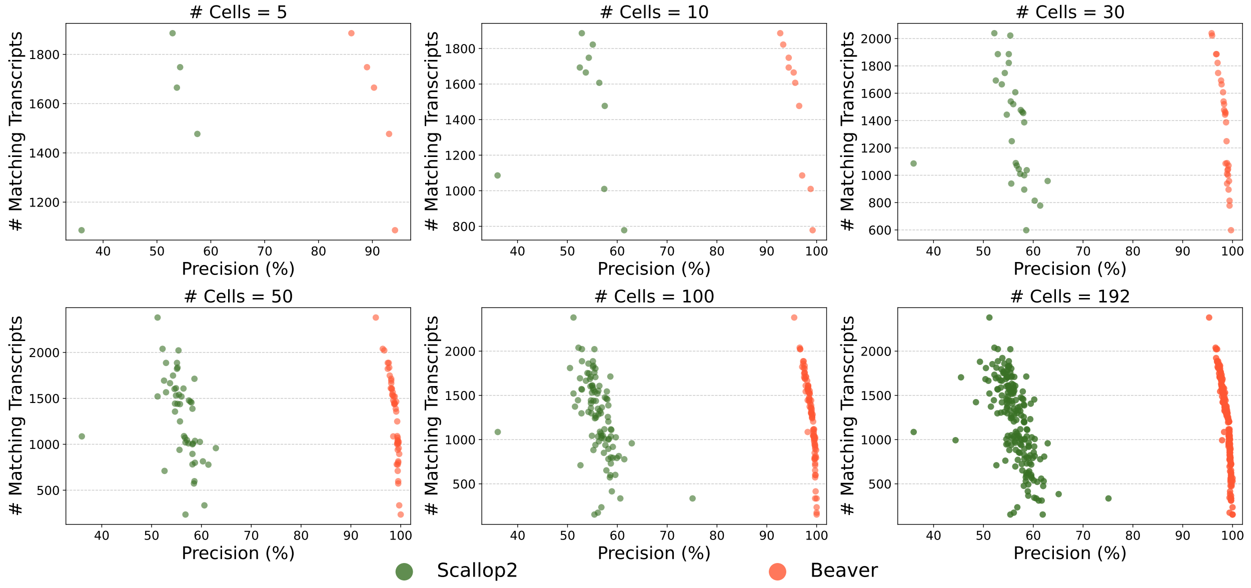


**Supplementary Figure 5**: Comparison of adjusted precision between Beaver and Scallop2 across varying cell population sizes ($n = 5, 10, 30, 50, 100, 192$) in the HEK293T dataset. Each data point represents an individual cell.

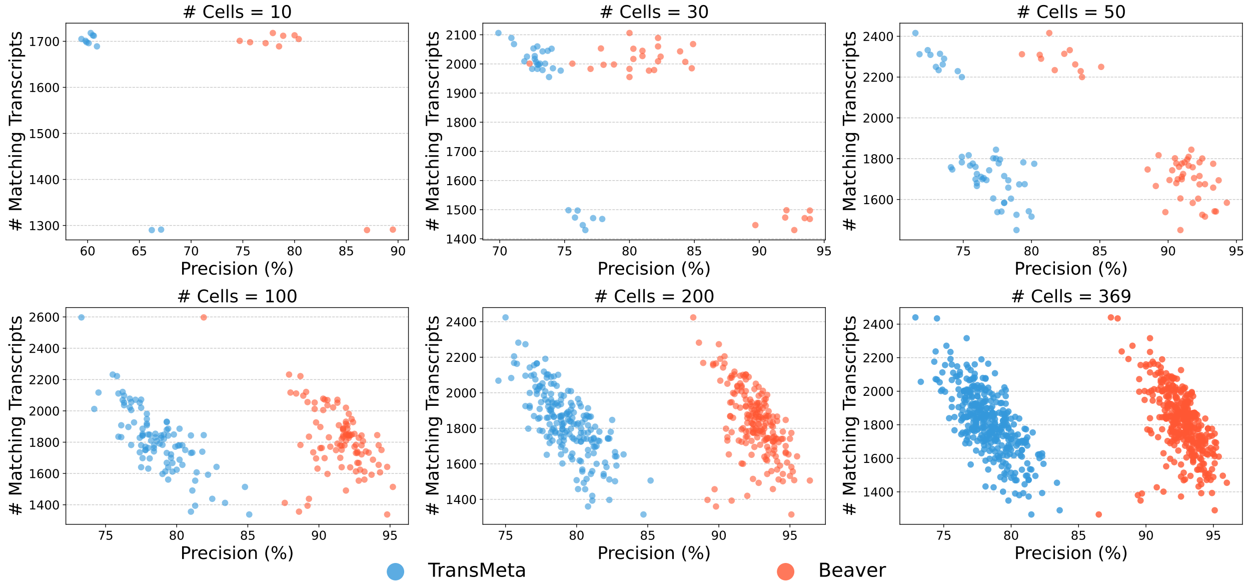


**Supplementary Figure 6**: Comparison of adjusted precision between Beaver and TransMeta across varying cell population sizes ($n = 10, 30, 50, 100, 200, 369$) in the Mouse-Fibroblast dataset. Each data point represents an individual cell.

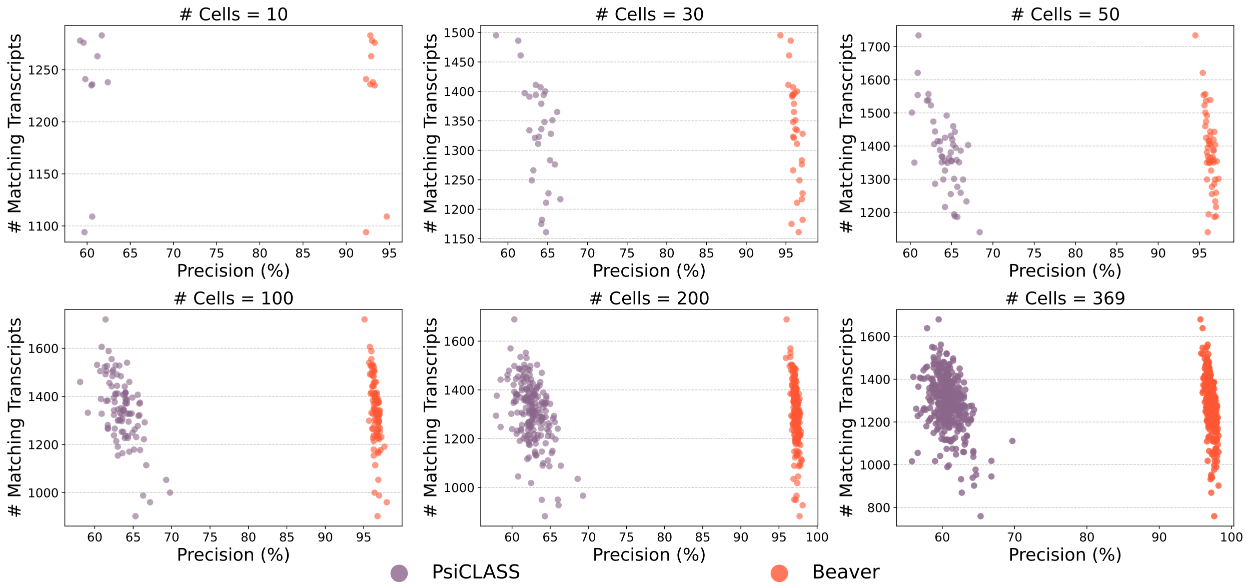


**Supplementary Figure 7**: Comparison of adjusted precision between Beaver and PsiCLASS across varying cell population sizes ($n = 10, 30, 50, 100, 200, 369$) in the Mouse-Fibroblast dataset. Each data point represents an individual cell.

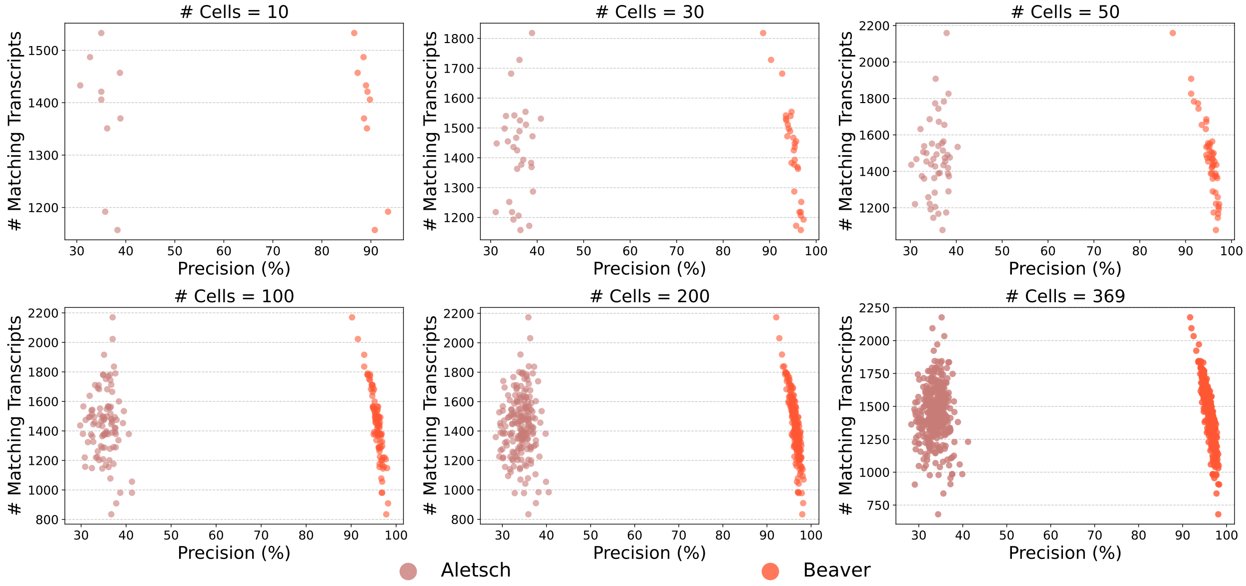


**Supplementary Figure 8**: Comparison of adjusted precision between Beaver and Aletsch across varying cell population sizes ($n = 10, 30, 50, 100, 200, 369$) in the Mouse-Fibroblast dataset. Each data point represents an individual cell.

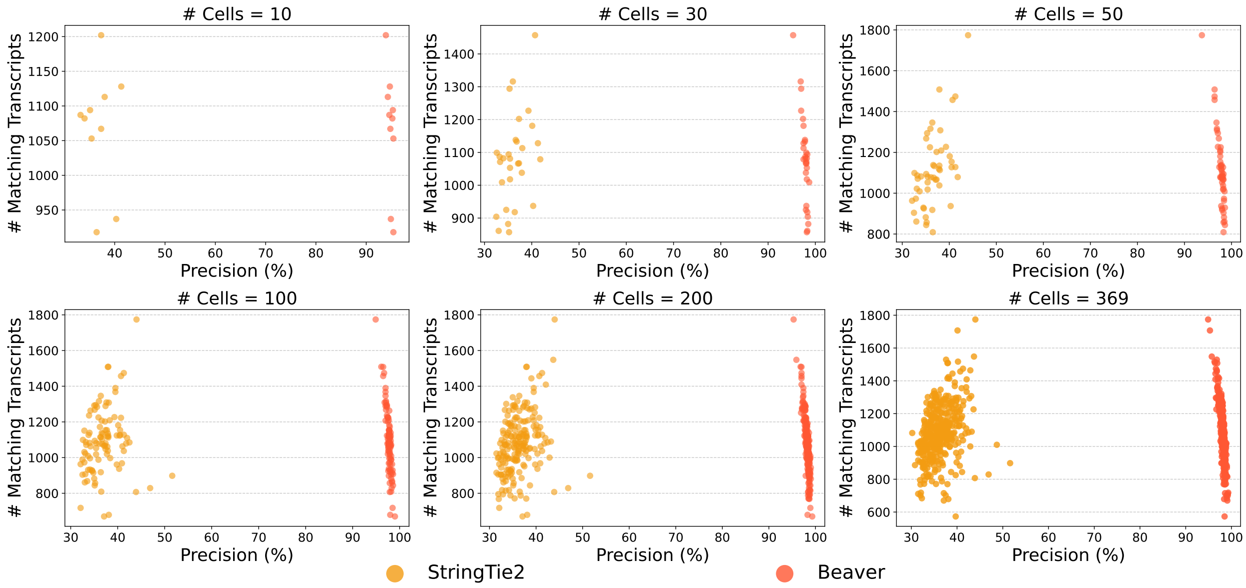


**Supplementary Figure 9**: Comparison of adjusted precision between Beaver and StringTie2 across varying cell population sizes ($n = 10, 30, 50, 100, 200, 369$) in the Mouse-Fibroblast dataset. Each data point represents an individual cell.

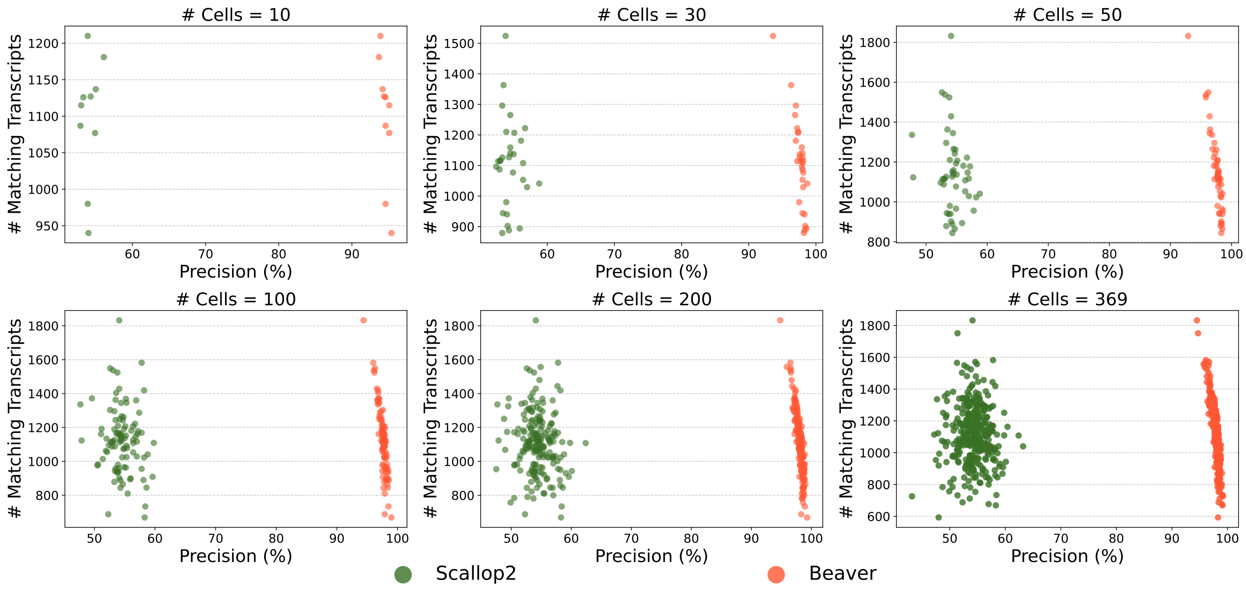


**Supplementary Figure 10**: Comparison of adjusted precision between Beaver and Scallop2 across varying cell population sizes ($n = 10, 30, 50, 100, 200, 369$) in the Mouse-Fibroblast dataset. Each data point represents an individual cell.

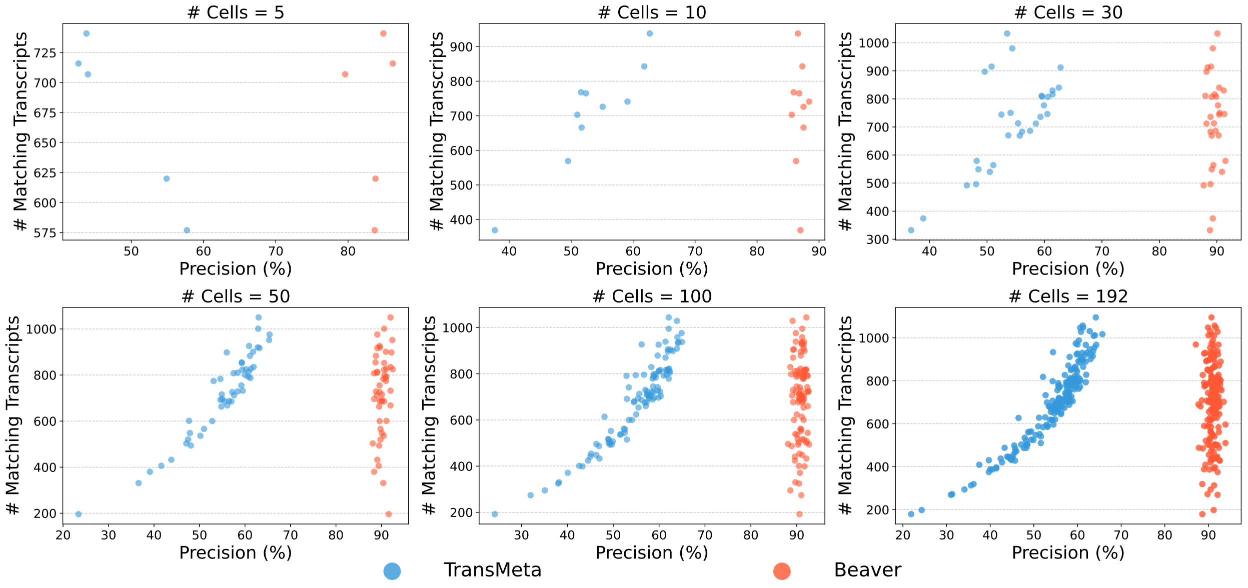


**Supplementary Figure 11**: Comparison of adjusted precision between Beaver and TransMeta across varying cell population sizes ($n = 5, 10, 30, 50, 100, 192$) in the simulated HEK293T-Sim dataset. Each data point represents an individual cell.

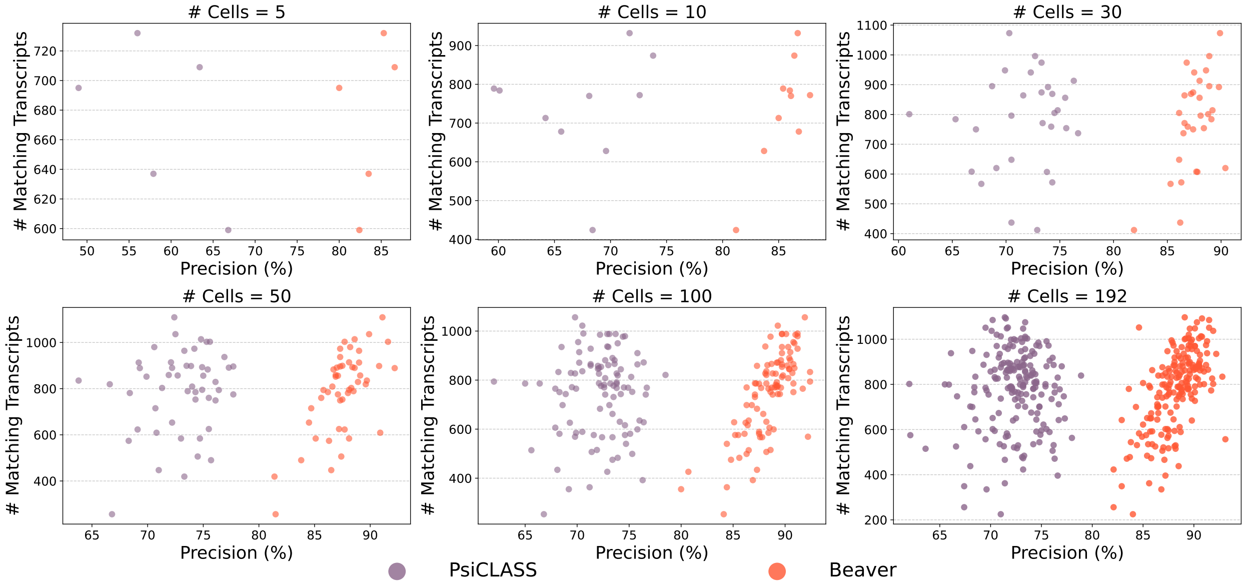


**Supplementary Figure 12**: Comparison of adjusted precision between Beaver and PsiCLASS across varying cell population sizes ($n = 5, 10, 30, 50, 100, 192$) in the simulated HEK293T-Sim dataset. Each data point represents an individual cell.

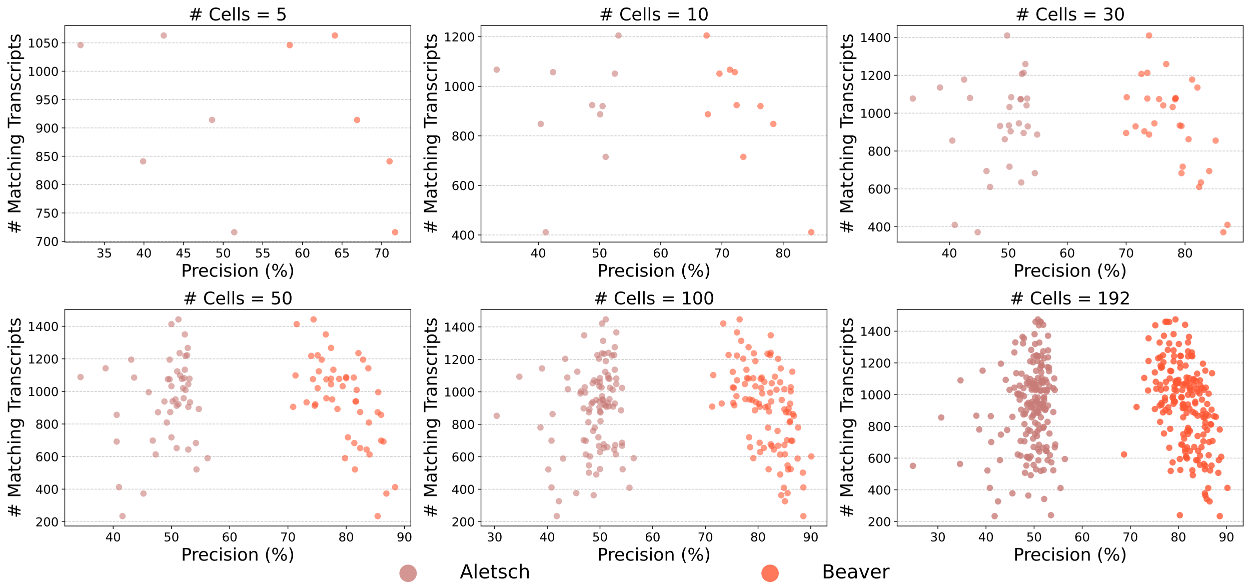


**Supplementary Figure 13**: Comparison of adjusted precision between Beaver and Aletsch across varying cell population sizes ($n = 5, 10, 30, 50, 100, 192$) in the simulated HEK293T-Sim dataset. Each data point represents an individual cell.

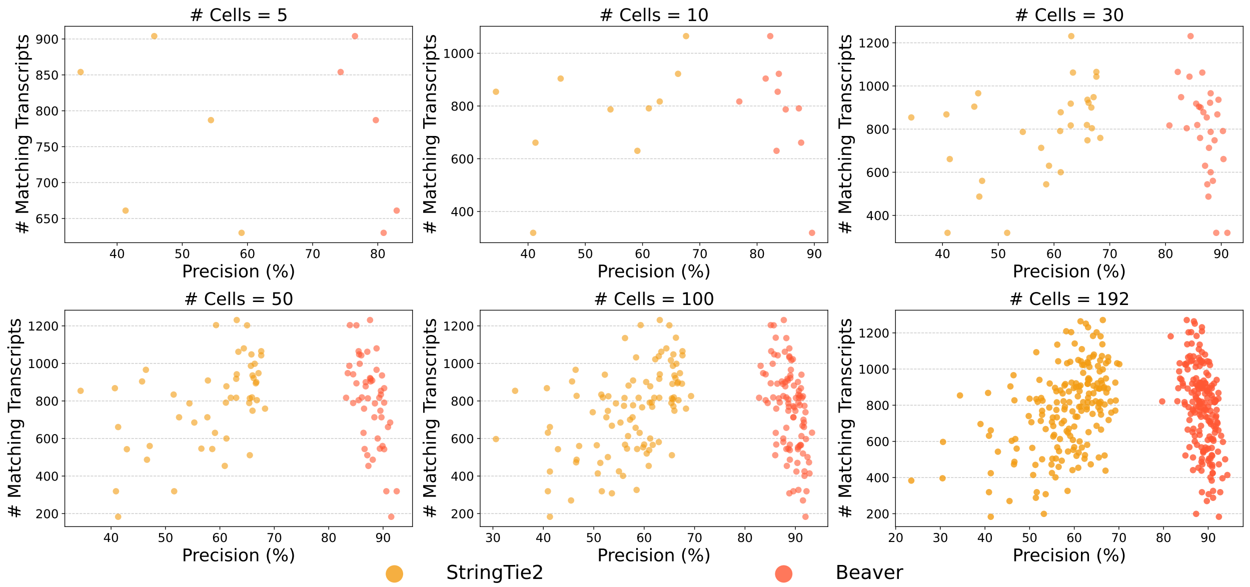


**Supplementary Figure 14**: Comparison of adjusted precision between Beaver and StringTie2 across varying cell population sizes ($n = 5, 10, 30, 50, 100, 192$) in the simulated HEK293T-Sim dataset. Each data point represents an individual cell.

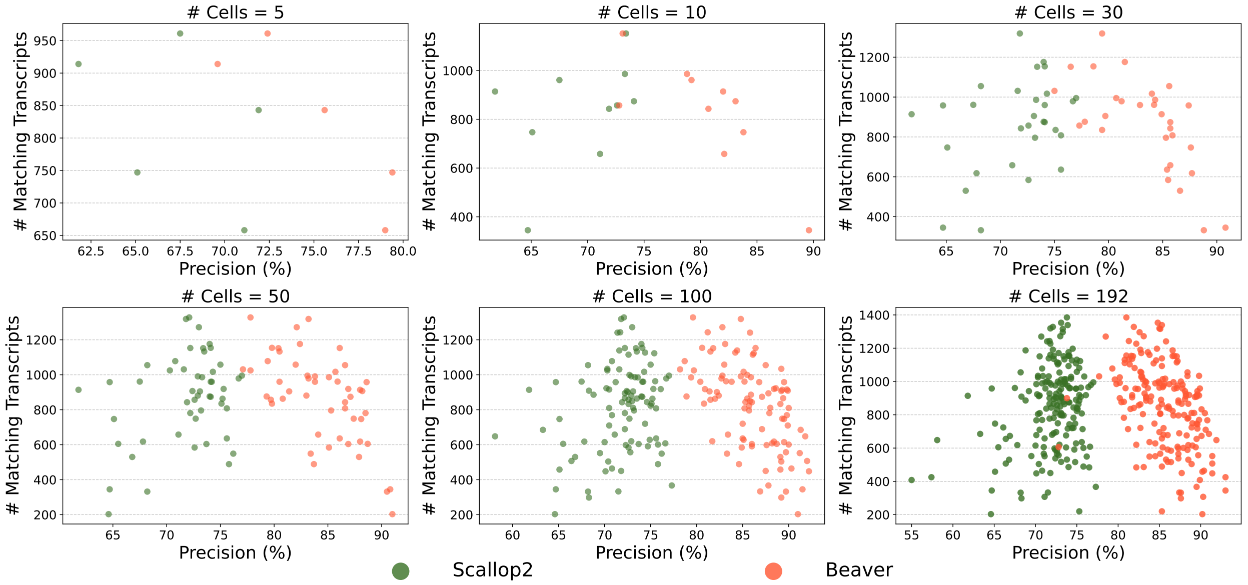


**Supplementary Figure 15**: Comparison of adjusted precision between Beaver and Scallop2 across varying cell population sizes ($n = 5, 10, 30, 50, 100, 192$) in the simulated HEK293T-Sim dataset. Each data point represents an individual cell.

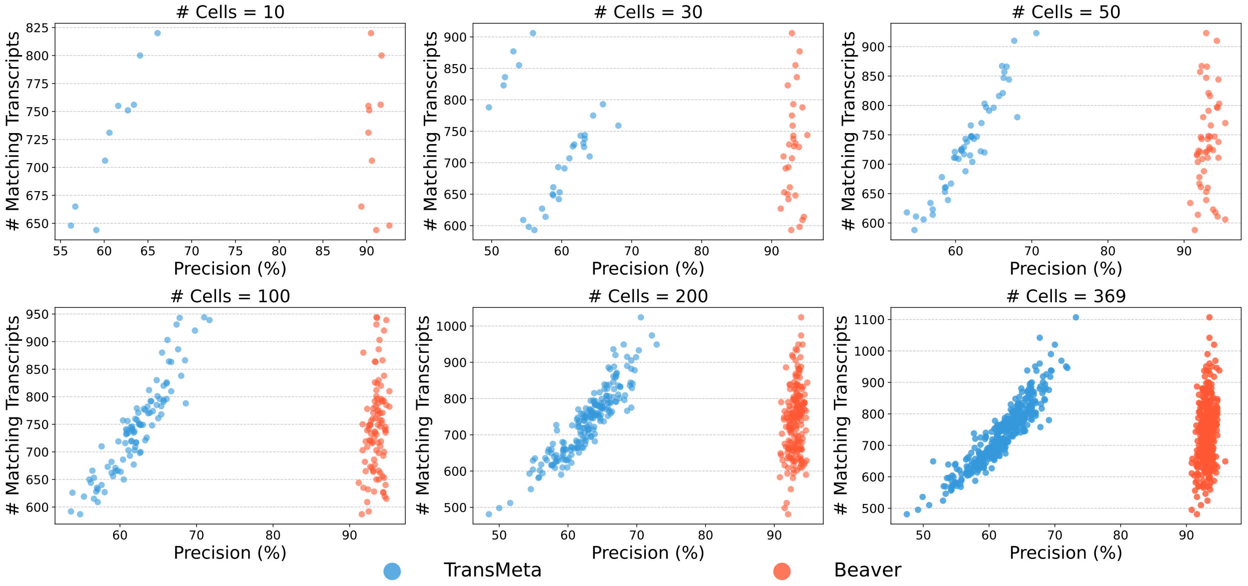


**Supplementary Figure 16**: Comparison of adjusted precision between Beaver and TransMeta across varying cell population sizes ($n = 10, 30, 50, 100, 200, 369$) in the simulated Mouse-Fibroblast-Sim dataset. Each data point represents an individual cell.

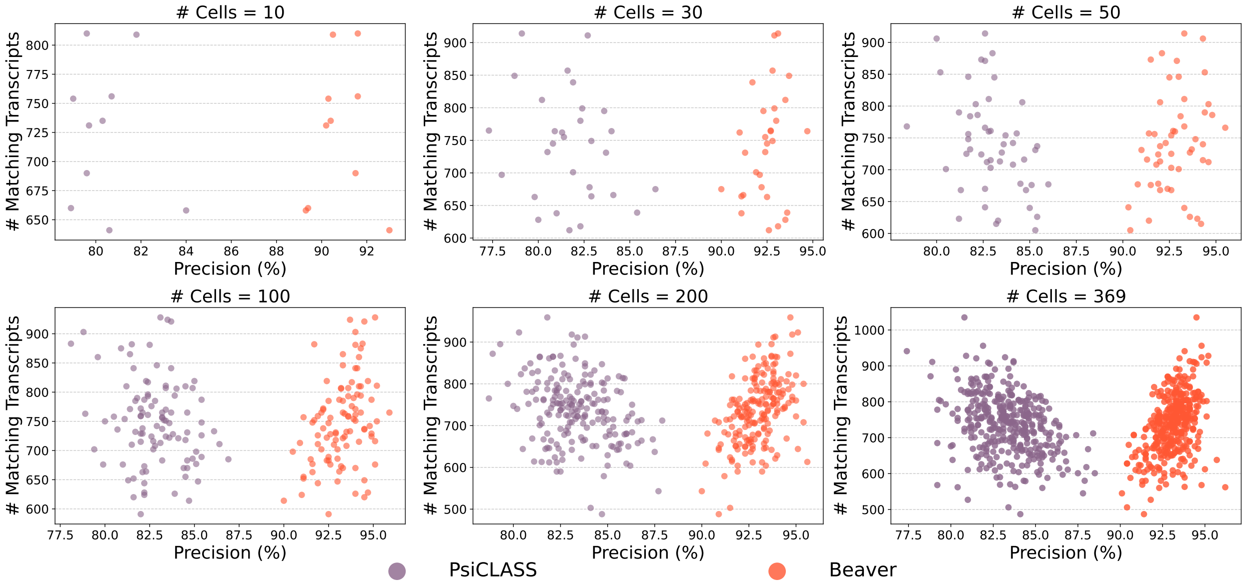


**Supplementary Figure 17**: Comparison of adjusted precision between Beaver and PsiCLASS across varying cell population sizes ($n = 10, 30, 50, 100, 200, 369$) in the simulated Mouse-Fibroblast-Sim dataset. Each data point represents an individual cell.

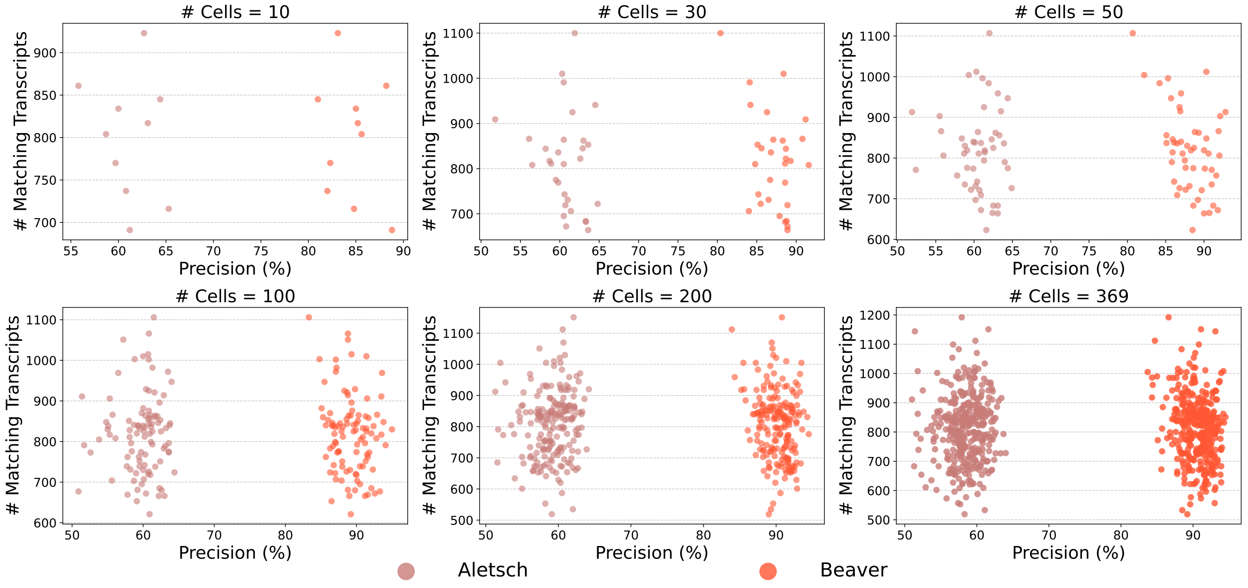


**Supplementary Figure 18**: Comparison of adjusted precision between Beaver and Aletsch across varying cell population sizes ($n = 10, 30, 50, 100, 200, 369$) in the simulated Mouse-Fibroblast-Sim dataset. Each data point represents an individual cell.

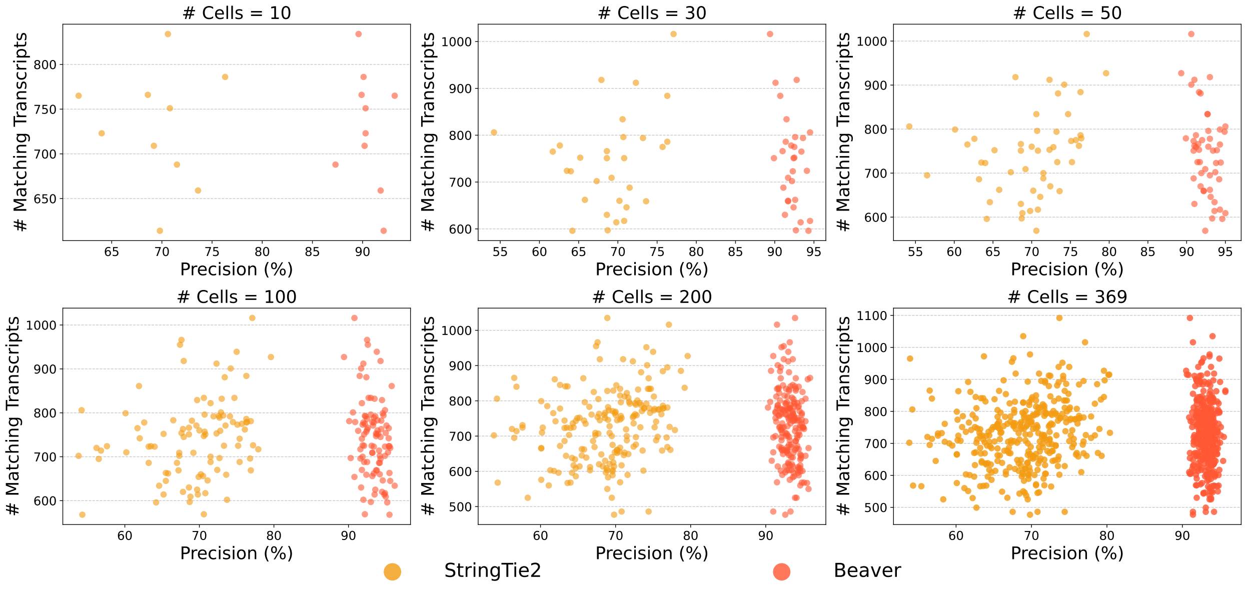


**Supplementary Figure 19**: Comparison of adjusted precision between Beaver and StringTie2 across varying cell population sizes ($n = 10, 30, 50, 100, 200, 369$) in the simulated Mouse-Fibroblast-Sim dataset. Each data point represents an individual cell.

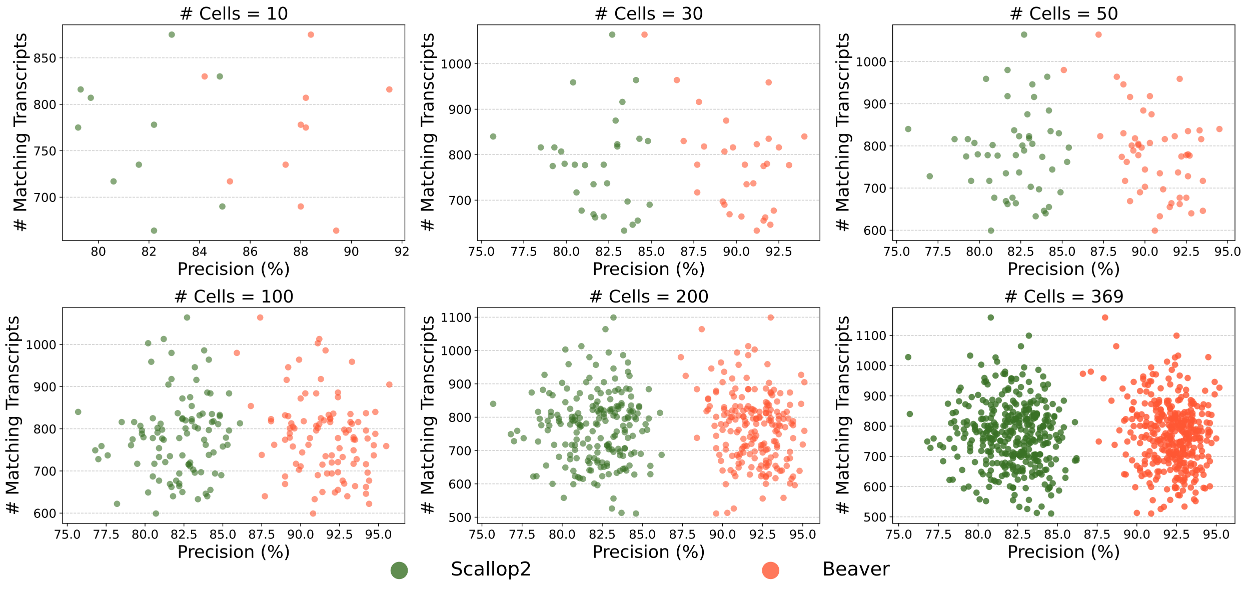


**Supplementary Figure 20**: Comparison of adjusted precision between Beaver and Scallop2 across varying cell population sizes ($n = 10, 30, 50, 100, 200, 369$) in the simulated Mouse-Fibroblast-Sim dataset. Each data point represents an individual cell.
